# *Arabidopsis* UGT76B1 glycosylates *N*-hydroxy-pipecolic acid and inactivates systemic acquired resistance in tomato

**DOI:** 10.1101/2020.07.06.189894

**Authors:** Eric C. Holmes, Yun-Chu Chen, Mary Beth Mudgett, Elizabeth S. Sattely

**Affiliations:** Department of Chemical Engineering, Stanford University, Stanford, CA, 94305, USA; Department of Biology, Stanford University, Stanford, CA, 94305, USA; Howard Hughes Medical Institute

## Abstract

Systemic acquired resistance (SAR) is a mechanism that plants utilize to connect a local pathogen infection to global defense responses. *N*-hydroxy-pipecolic acid (NHP) and a glycosylated derivative are produced during SAR, yet their individual roles in the response have not yet been elucidated. Here we report that *Arabidopsis thaliana* UGT76B1 can generate glycosylated NHP (NHP-Glc) *in vitro* and when transiently expressed alongside Arabidopsis NHP biosynthetic genes in two Solanaceous plants. During infection, Arabidopsis *ugt76b1* mutants do not accumulate NHP-Glc and accumulate less glycosylated salicylic acid (SA-Glc) than wild type plants. The metabolic changes in *ugt76b1* mutant plants are accompanied by enhanced defense to the bacterial pathogen *Pseudomonas syringae*, suggesting that glycosylation of SAR molecules NHP and SA by UGT76B1 plays an important role in defense modulation. Transient expression of Arabidopsis *UGT76B1* with the Arabidopsis NHP biosynthesis genes *ALD1* and *FMO1* in tomato increases NHP-Glc production and reduces NHP accumulation in local tissue, and abolishes the systemic resistance seen when expressing NHP-biosynthetic genes alone. These findings reveal that the glycosylation of NHP by UGT76B1 alters defense priming in systemic tissue and provide further evidence for the role of the NHP aglycone as the active metabolite in SAR signaling.

## Introduction

Systemic acquired immunity in plants is a coordinated defense response that leads to heightened disease protection throughout the plant body following an initial, localized pathogen attack. Several small molecules have been found to help orchestrate this process, including the ubiquitous hormone salicylic acid (SA) (Klessig et al., 2018), and the newly-discovered the signaling metabolite *N*-hydroxy-pipecolic acid (NHP) (Chen et al., 2018; Hartmann et al., 2018), which is thought to have a lead role in SAR. The enzyme flavin monooxygenase 1 (FMO1) catalyzes the *N*-hydroxylation of the nonproteinogenic amino acid pipecolic acid (Pip) in the biosynthesis of NHP and is required for the initiation and amplification to SAR signaling (Chen et al., 2018; Hartmann et al., 2018).

Both SA and NHP can be isolated from plants with several metabolic modifications, most notably as the glycosylated derivatives. In prior work using Arabidopsis plants, we and others have observed that both NHP and its hexose-conjugated derivative (NHP-Glc) accumulate after bacterial infection in seedlings and leaves (Chen et al., 2018; Hartmann and Zeier, 2018). Both NHP-Glc and the aglycone are absent from unelicited plants and pathway mutants deficient in FMO1, prompting questions about the role of NHP glycosylation in SAR. The glycosyltransferase required for generation of NHP-Glc however has remained elusive.

Structural modifications of small plant signaling molecules appear to have evolved as a dynamic mechanism to modulate the activity of these chemical signals. Common enzymatic modifications to base hormone scaffolds include hydroxylation, carboxylation, sulfation, acetylation, methylation, amino acid conjugation, and glycosylation (Westfall et al., 2013). Some hormones, such as the defense hormone jasmonic acid, undergo multiple enzymatic modifications (Wasternack and Hause, 2013) to create bioactive (Staswick and Tiryaki, 2004), inactive (Smirnova et al., 2017), and differentially active (Nakamura et al., 2011) compounds. Often, loss of function mutants of these modifying enzymes can have severe impact on plant physiology, leading to developmental phenotypes in the case of auxins (Nakazawa et al., 2001; Takase et al., 2004; Staswick et al., 2005), brassinosteroids (Choi et al., 2013), and gibberellins (Wang et al., 2012) and to altered responses to environmental stresses in the case of abscisic acid (Liu et al., 2015), jasmonic acid (JA) (Caarls et al., 2017; Smirnova et al., 2017), and SA (Liu et al., 2009; Boachon et al., 2014). In some instances, hormone conjugation appears to serve as a reservoir of a molecule for fast deployment, in other cases it seems to be a metabolic mechanism for attenuating activity and depleting the active form (Piotrowska and Bajguz, 2011).

Several lines of evidence show that the NHP aglycone is sufficient to initiate SAR signaling but have not yet revealed a functional role for glycosylation. For example, the treatment of *Arabidopsis thaliana* (Chen et al., 2018; Hartmann et al., 2018), *Capsicum annuum* (sweet pepper) (Holmes et al., 2019), or *Solanum lycopersicum* (tomato) (Holmes et al., 2019) leaves with synthetic NHP induces resistance against bacterial infection in distal tissues not treated with NHP. Furthermore, transient overexpression of the Arabidopsis NHP biosynthetic enzymes AGD2-like defense protein 1 (ALD1; (Navarova et al., 2012)) and FMO1 (Chen et al., 2018; Hartmann et al., 2018) leads to the production of NHP in tomato leaves and results in enhanced resistance to bacterial infection in distal tissues (Holmes et al., 2019). Notably, NHP-Glc was not detected in the NHP treated tomato leaves, suggesting that NHP-Glc synthesis and/or accumulation may not occur in tomato. These data, coupled with the observation that NHP-Glc does not accumulate in the absence of infection in Arabidopsis, suggests NHP-Glc is not simply a storage form of NHP.

Despite the clear role of NHP biosynthesis for the initiation of systemic resistance, several open questions remain regarding (i) the active form of NHP metabolites, (ii) the potential role of NHP glycosylation in modulating SAR signaling, and (iii) more broadly, mechanisms of signal initiation, transport, and attenuation in plant systemic resistance. In an effort to better understand the potential role of NHP-Glc in the SAR response, we sought to establish the genetic and biochemical basis for NHP glycosylation in Arabidopsis and test the influence of the putative glycosylating enzyme(s) in the SAR-mediated disease resistance. Here we report that Arabidopsis UDP-glycosyltransferase UGT76B1 can generate glycosylated NHP (NHP-Glc) *in vitro* and when transiently expressed alongside Arabidopsis NHP biosynthetic genes in two Solanaceous plants.

Our results provide new insight into how plants use specific metabolic transformations to alter the behavior of the key signaling molecule NHP in systemic defense.

## Results

### Heterologously expressed Arabidopsis UGT76B1 glycosylates NHP in planta

Previous studies have indicated that NHP-Glc accumulates in Arabidopsis after pathogen infection (Chen et al., 2018; Hartmann and Zeier, 2018). We hypothesized that a dedicated NHP-glycosyltransferase may be highly expressed under pathogen stress conditions (Figure 1A). We analyzed a set of publicly available microarray datasets for the mRNA expression pattern of 103 Arabidopsis UDP-dependent glycosyltransferase genes (*UGTs*) under various biotic stress conditions (Supplementary Figure 1). We prioritized testing of candidate *UGTs* based upon their high level of mRNA abundance across all biotic stress conditions and selected a few others based upon their high level of mRNA abundance under a specific pathogen stress. For the initial screen, we selected 14 *UGTs* from this microarray analysis (Supplementary Figure 1) as well as four additional *UGTs* (*UGT73B2*, *UGT73B3*, *UGT73C3*, and *UGT73C5*) based on expression profiles in RNA sequencing experiments (Bernsdorff et al., 2016; Hartmann et al., 2018). Our goal was to first identify Arabidopsis *UGT*s that could generate NHP during heterologous expression and subsequently determine the role of any candidates in Arabidopsis.

**Figure 1.**
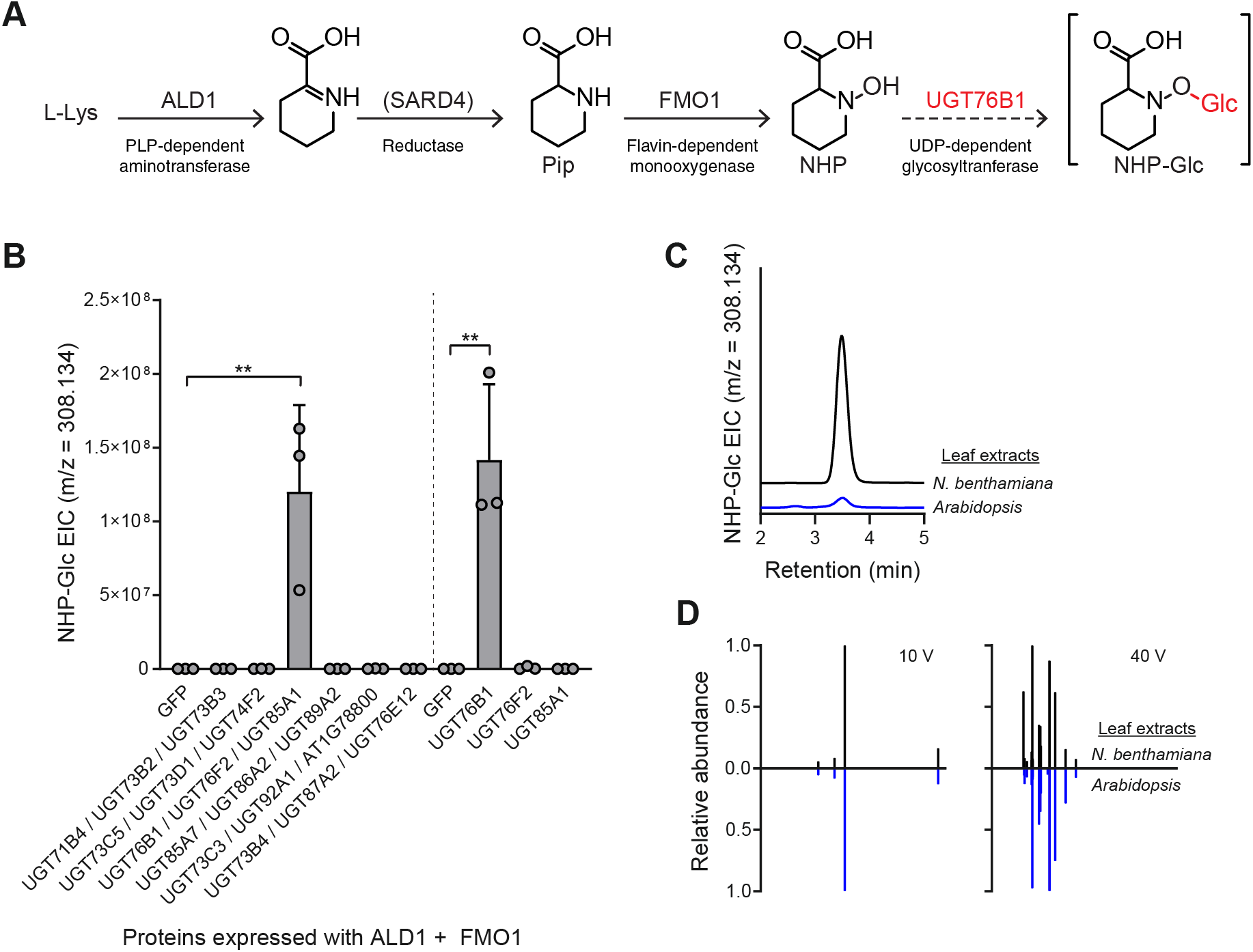
Screen of 18 Arabidopsis UGTs for their ability to glycosylate NHP. (A) Biosynthetic pathway for production of NHP-Glc from L-Lys in Arabidopsis. The biosynthetic activity of UGT76B1 was characterized in this work. (B) Abundance of NHP-Glc measured with LC-MS after transient expression of GFP or respective Arabidopsis UGTs alongside Arabidopsis ALD1 + FMO1 in *N. benthamiana* leaves. In initial screen, *Agrobacteria* strains harboring distinct UGTs were combined in equal proportions and co-infiltrated with *Agrobacteria* strains harboring *ALD1* and *FMO1*. In second screen, *Agrobacteria* strains harboring *UGT76B1*, *UGT76F2*, or *UGT85A1* were separately co-infiltrated with *Agrobacteria* strains harboring *ALD1* and *FMO1*. Total inoculum (OD_600_) was kept constant in both experiments by including *Agrobacteria* harboring *GFP* as a control. Bars represent the mean ± SD (*n* = 3 independent biological replicates). Values reported as zero indicate no detection of metabolites. Asterisks indicate a significant NHP-Glc increase (one-tailed *t* test; \*\**P* < 0.01). (C) Representative LC-MS chromatograms of NHP-Glc (m/z = 308.134) in extracts from transient expression of ALD1 + FMO1 + UGT76B1 in *N. benthamiana* (black) and Arabidopsis adult leaves (blue) after infiltration with 1 mM NHP synthetic standard. (D) Comparative MS/MS spectra of NHP-Glc in extracts from transient expression of ALD1 + FMO1 + UGT76B1 in *N. benthamiana* (black) and Arabidopsis adult leaves (blue) after infiltration with 1 mM NHP synthetic standard at collision energies of 10V and 40V.

In previous studies, we used *Agrobacterium*-mediated transient expression in *Nicotiana benthamiana* (Kapila et al., 1997) as a heterologous expression platform to produce NHP *in planta* (Chen et al., 2018; Holmes et al., 2019). Under these transient expression conditions, NHP-Glc was not detected in extracts from *N. benthamiana* leaves (Chen et al., 2018; Holmes et al., 2019). We hypothesized that this heterologous expression system could be used to screen Arabidopsis UGT candidates with minimal background signal from native enzymes. We cloned 18 candidate *UGT* cDNAs and then transiently expressed them with Arabidopsis *ALD1* and *FMO1* (the minimal set of genes required for NHP biosynthesis (Holmes et al., 2019) in *N. benthamiana* leaves. To expedite testing and metabolite analysis, we expressed our candidate enzymes in groups of three by combining *Agrobacteria* strains harboring separate *UGT* candidates and *GFP* in equal proportions and co-infiltrated them with *Agrobacteria* harboring *ALD1* and *FMO1*.

Liquid chromatography-mass spectrometry (LC-MS) analysis of methanolic extracts from these leaves revealed that one set of genes tested (*UGT76B1*, *UGT76F2*, and *UGT85A1*) led to significant accumulation of NHP-Glc when coexpressed with *ALD1* and *FMO1* compared to coexpression of *ALD1* and *FMO1* with *GFP* (Figure 1B). We then transiently expressed each of these respective *UGTs* with *ALD1* and *FMO1* and found that leaves expressing *UGT76B1* (At3g11340) were the only ones that accumulated a significant amount of NHP-Glc (Figure 1B). *N. benthamiana* leaves transiently expressing *ALD1*, *FMO1*, and *UGT76B1* accumulated significantly less free NHP (as measured using LC-MS) than did leaves expressing *ALD1* and *FMO1* alone, indicating a high conversion rate from NHP to NHP-Glc by UGT76B1 (Supplementary Figure 2). The compound produced in *N. benthamiana* had the same LC-MS retention time (Figure 1C) and MS/MS fragmentation pattern (Figure 1D) as NHP-Glc produced in adult Arabidopsis leaves, suggesting that Arabidopsis UGT76B1 is producing the same glycosylated NHP derivative as natively accumulates in Arabidopsis.

### In vitro biochemistry of UGT76B1 expressed from N. benthamiana and E. coli

Previous studies have shown that UGT76B1 glycosylates the plant hormone salicylic acid (SA) and the isoleucine catabolite 2-hydroxy-3-methyl-pentanoic acid (ILA) *in vitro* and contributes to the accumulation of their respective glycosides *in planta* (von Saint Paul et al., 2011; Noutoshi et al., 2012; Maksym et al., 2018; Bauer et al., 2020). Our results in *N. benthamiana* suggested that UGT76B1 could glycosylate a third defense-related metabolite, NHP. To confirm these previous results and the determine that the NHP-Glc we detected in *N. benthamiana* was a direct result of UGT76B1 activity, we spiked crude protein extracts from *N. benthamiana* leaves expressing GFP or UGT76B1 with UDP-glucose and the aglycone substrates ILA, SA, or NHP. Protein extracts from leaves expressing GFP did not produce any of the respective glycosides while extracts from leaves expressing UGT76B1 produced all three (Figure 2A). Furthermore, we transiently expressed His-tagged UGT76B1 in *N. benthamiana* leaves and enriched for UGT76B1 using Ni-NTA affinity purification. Partially purified UGT76B1-6xHis from *N. benthamiana* catalyzed the synthesis of NHP-Glc *in vitro* while denatured protein did not (Figure 2B).

**Figure 2.**
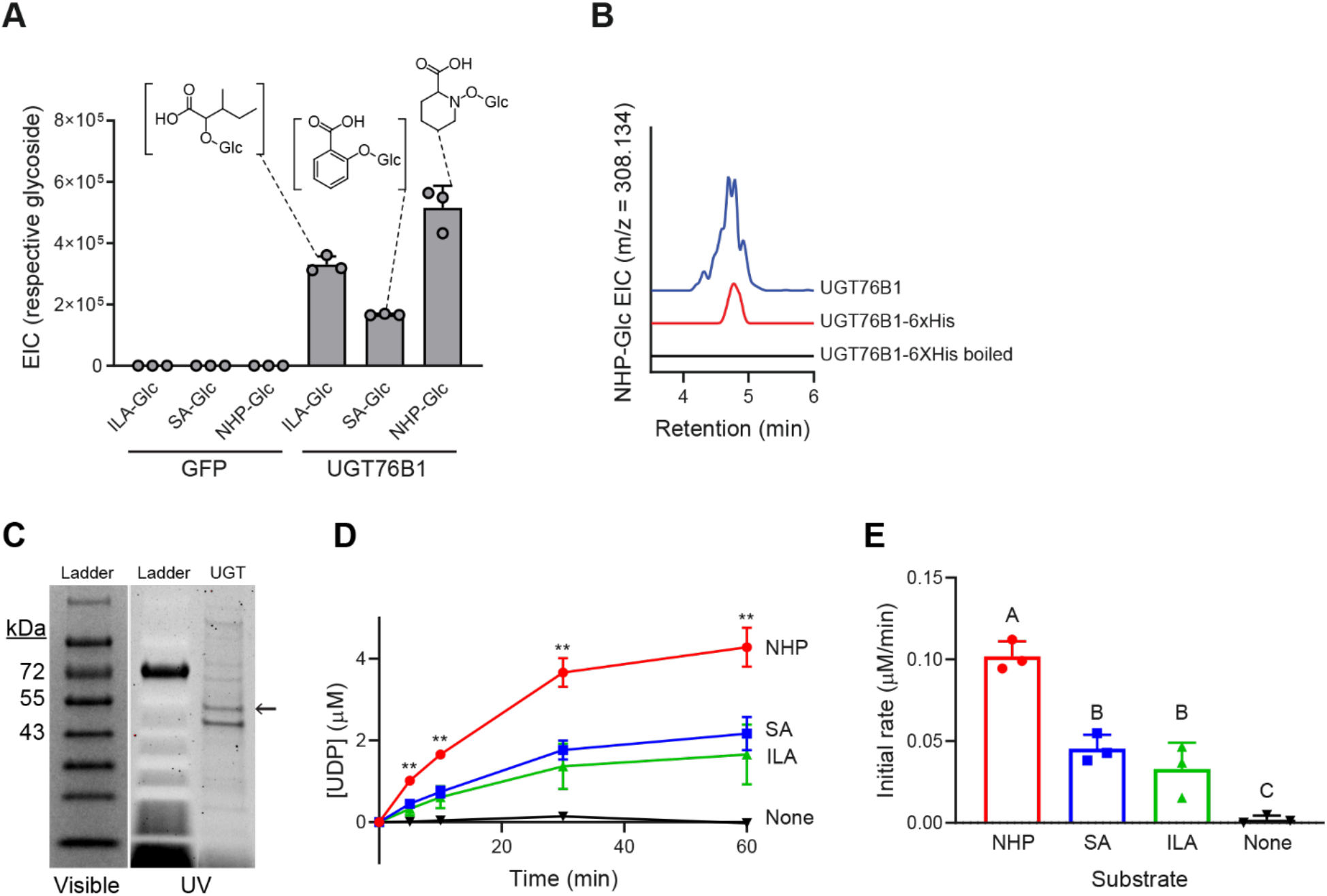
*In vitro* characterization of Arabidopsis UGT76B1. (A) GFP or Arabidopsis UGT76B1 were transiently expressed in *N. benthamiana* leaves and crude protein extracts were incubated with 5 mM UDP glucose and 1 mM aglycone substrates (2-hydroxy-3-methylvaleric acid (ILA), salicylic acid (SA), or NHP). Levels indicate abundances of glycosides measured with LC-MS after 3 h incubation. Bars represent the mean ± SD (*n* = 3 independent biological replicates). m/z used for quantification are: ILA-Glc ([M-H]^−^ = 293.124), SA-Glc ([M-H]^−^ = 299.077), and NHP-Glc ([M+H]^+^ = 308.134). (B) Representative LC-MS chromatograms of NHP-Glc (m/z = 308.134) from crude extract from *N. benthamiana* leaves transiently expressing Arabidopsis UGT76B1 (blue), and enriched (red) or denatured (black) UGT76B1-6xHis purified from *N. benthamiana* leaves. (C) SDS-PAGE gel of enriched UGT76B1-6xHis purified from *E. coli*. Same gel imaged under visible light and UV light is included to better visualize ladder bands. Expected mass of UGT76B1-6xHis is ~51 kDa. (D) Enriched UGT76B1-6xHis from *E. coli* was incubated with NHP (red), SA (blue), ILA (green), or no substrate (black) *in vitro*. Aliquots were quenched at increasing time points and free UDP liberated from the reaction of UGT76B1 with its respective substrates was measured using an enzyme-linked assay. Asterisks indicate a significant difference (two-tailed *t* test; \*\**P* < 0.01). Points represent the mean ± SD (*n* = 3 independent biological replicates). (E) Initial rate of reaction from (D) was measured as the slope from t = 0 to t = 5 min. Letters indicate significantly different groups using two-tailed t-tests (*P* < 0.01). Bars represent the mean ± SD (*n* = 3 independent biological replicates).

Given the promiscuity of some plant UGTs on structurally-similar substrates (Lim et al., 2002), it is unsurprising that UGT76B1 can glycosylate ILA, SA, and NHP. To better understand the ability of this enzyme to glycosylate these substrates, we expressed and enriched UGT76B1-6xHis from *E. coli* (Figure 2C) and then tested its activity using an enzyme-coupled assay (Zegzouti et al., 2013). During each UGT catalytic reaction, UDP is released and the concentration of free UDP in a given reaction can be directly measured using this assay. Reactions with NHP generated significantly more UDP over the course of one hour than was generated with SA or ILA as substrates (Figure 2D). The initial rate of reaction with NHP was also approximately 2x faster than with either SA or ILA (Figure 2E). These results confirm that UGT76B1 acts on ILA, SA, and NHP and indicates it is more active on NHP as a substrate in these conditions.

To determine where glucose conjugation is occurring on NHP, we derivatized synthetic NHP and an extract from *N. benthamiana* leaves transiently expressing ALD, FMO1, and UGT76B1 with trimethylsilyldiazomethane (TMSD), a reagent commonly used to selectively methylate carboxylic acids (Kühnel et al., 2007; Topolewska et al., 2015). Derivatization of synthetic NHP generated a major, singly methylated product and a minor doubly methylated product (Supplementary Figure 3A). We hypothesize the singly methylated product to be NHP methyl ester based upon MS/MS fragmentation and the reported activity of TMSD (Supplementary Figure 3A). Derivatization of the *N. benthamiana* extract revealed a methylated NHP-Glc product with an MS/MS fragmentation pattern that matches that of NHP methyl ester (Supplementary Figure 3B), suggesting that UGT76B1 is generating NHP-β-D glucoside. UGT76B1 is also known to generate the β-D glucoside of salicylic acid (SA-Glc) (von Saint Paul et al., 2011; Noutoshi et al., 2012). A synthetic standard of NHP-Glc (which is currently unavailable) is required to definitively elucidate the structure of the glycosylated NHP produced by UGT76B1.

### Arabidopsis ugt76b1 mutant plants are impaired in NHP-Glc and SA-Glc production

Given that UGT76B1 is capable of glycosylating NHP when expressed heterologously in *N. benthamiana*, we next sought to determine its native function in Arabidopsis. We obtained the Syngenta Arabidopsis Insertion Library (SAIL) (Sessions et al., 2002) T-DNA insertional line SAIL_1171_A11 (*ugt76b1-1*; furthermore *ugt76b1*) from the Arabidopsis Biological Resource Center (ABRC). This mutant line was previously used to study the function of *UGT76B1* (von Saint Paul et al., 2011). To quantify NHP and SA derivatives, we grew WT Arabidopsis Col-0 (furthermore WT) and *ugt76b1* plants axenically in hydroponic media for two weeks, treated seedlings with 10 mM MgCl_2_ (mock), *Pseudomonas syringae* pathovar tomato DC3000 (*Pst*), 1 mM NHP, or 100 μM SA, and then measured metabolites using GC-MS and LC-MS (Figure 3 and Supplementary Figure 4). WT plants had significantly higher abundance of NHP-Glc and SA-Glc than did *ugt76b1* in *Pst*-treated plants (Figure 3), highlighting an important contribution from UGT76B1 in the glycosylation of NHP and SA during infection. While NHP-treated *ugt76b1* plants did contain detectable NHP-Glc, the abundance was reduced over 99% when compared to WT plants, suggesting that UGT76B1 is the primary NHP glycosyltransferase in Arabidopsis (Supplemental Figure 4). There may be other minor enzymes that contribute to NHP glycosylation, but NHP-Glc was only detectable in *ugt76b1* plants when a high concentration of NHP was supplemented (Supplementary Figure 4) and not when treated with *Pst* (Figure 3). The abundance of SA-Glc was reduced approximately 60% in *ugt76b1* plants compared to WT when supplemented with SA (Supplementary Figure 4), suggesting that the activity of UGT76B1 may also contribute significantly to the glycosylation of SA in Arabidopsis.

**Figure 3.**
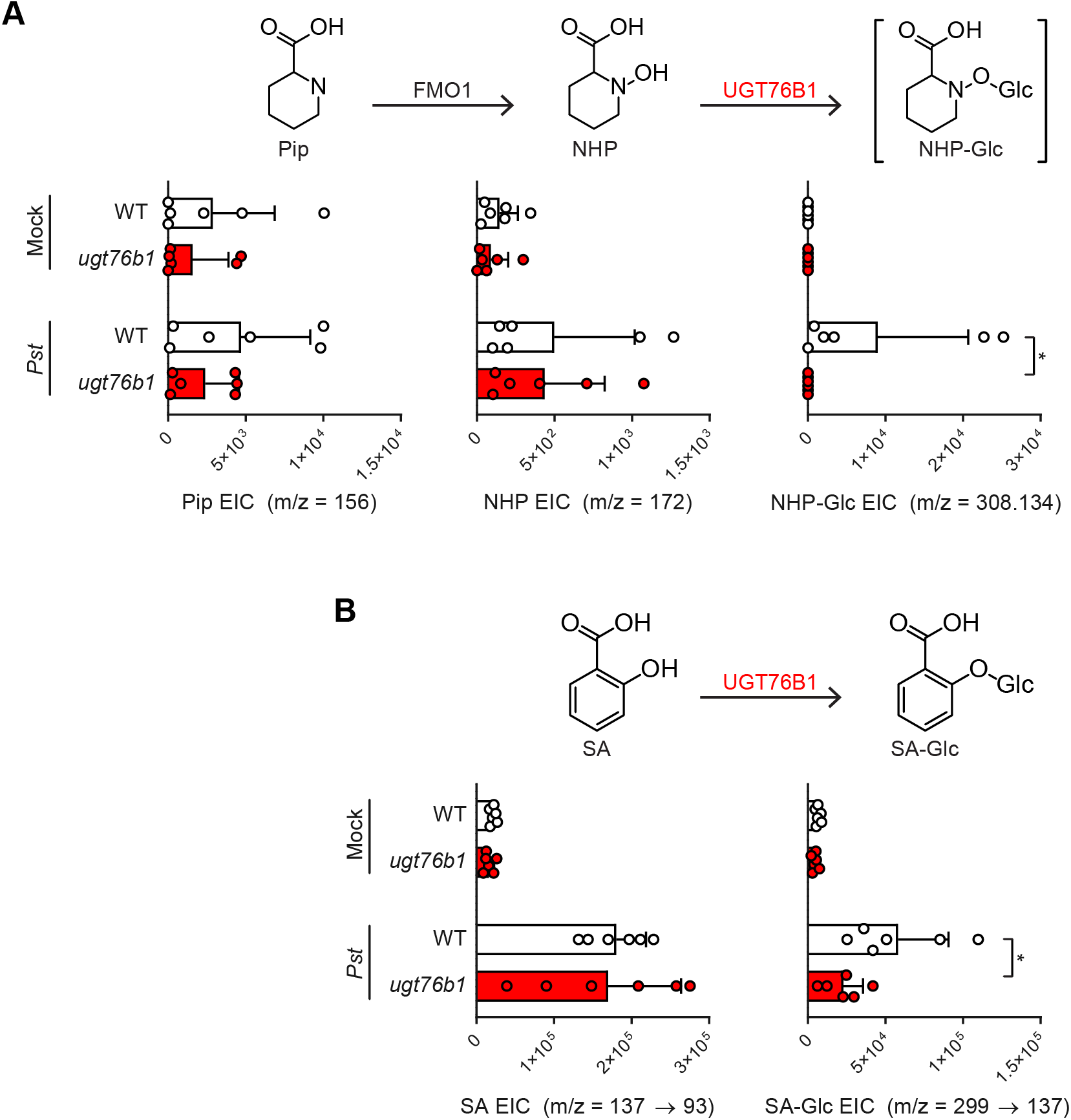
Abundance of NHP- and SA-related metabolites in WT and *ugt76b1* mutant seedlings. Arabidopsis WT (white bars) and *ugt76b1* (red bars) seedlings were grown axenically in hydroponic media for two weeks and treated with 10 mM MgCl_2_ (mock), or a suspension of *Pst* at OD_600_ of 0.01. After 24 h, seedlings were harvested and analyzed for NHP-related metabolites (A) and SA-related metabolites (B). Pip and NHP were measured as trimethylsilyl (TMS) and 2-TMS derivatives, respectively, using GC-MS. NHP-Glc, SA, and SA-Glc were measured using LC-MS. Bars represent the means ± SD (*n* = 6 independent biological replicates). Values reported as zero indicate no detection of metabolites. Asterisks indicate a significant metabolite decrease in *ugt76b1* plants (one-tailed *t* test; \**P* < 0.05).

### Arabidopsis ugt76b1 mutants are more resistant to bacterial infection

A previous study showed Arabidopsis *ugt76b1* mutants were more resistant to the biotrophic pathogen *Pst* and more susceptible to the necrotrophic pathogen *Alternaria brassicicola* (von Saint Paul et al., 2011), indicating that UGT76B1 plays a critical role in the regulation of disease resistance signaling. Given UGT76B1 can glycosylate NHP in Arabidopsis (Figure 3 and Supplemental Figure 4), we hypothesized that UGT76B1 may regulate the abundance of NHP that is available to initiate and sustain defense priming during SAR. To test this hypothesis and explore the function of NHP-Glc and UGT76B1, we performed SAR experiments as previously described (Chen et al., 2018; Hartmann et al., 2018). Briefly, three lower leaves (leaf number 5-7) of four-week-old WT, *ugt76b1*, and *fmo1* (a NHP and SAR deficient mutant) were infiltrated with 10 mM MgCl_2_ (mock) or a 5×10^6^ cfu/ml suspension of *Pst avrRpt2*, an avirulent strain that induces a strong defense response in WT plants. Two days later, an upper leaf of each plant was challenged with a 1×10^5^ cfu/ml suspension of *Psm* ES4326, a virulent strain (Figure 4A and Supplemental Figure 5A). Disease symptoms and titers of *Psm* ES4326 in the infected upper leaves were then photographed and quantified at 3 days post infiltration (dpi), respectively (Figure 4B and Supplemental Figure 5B).

**Figure 4.**
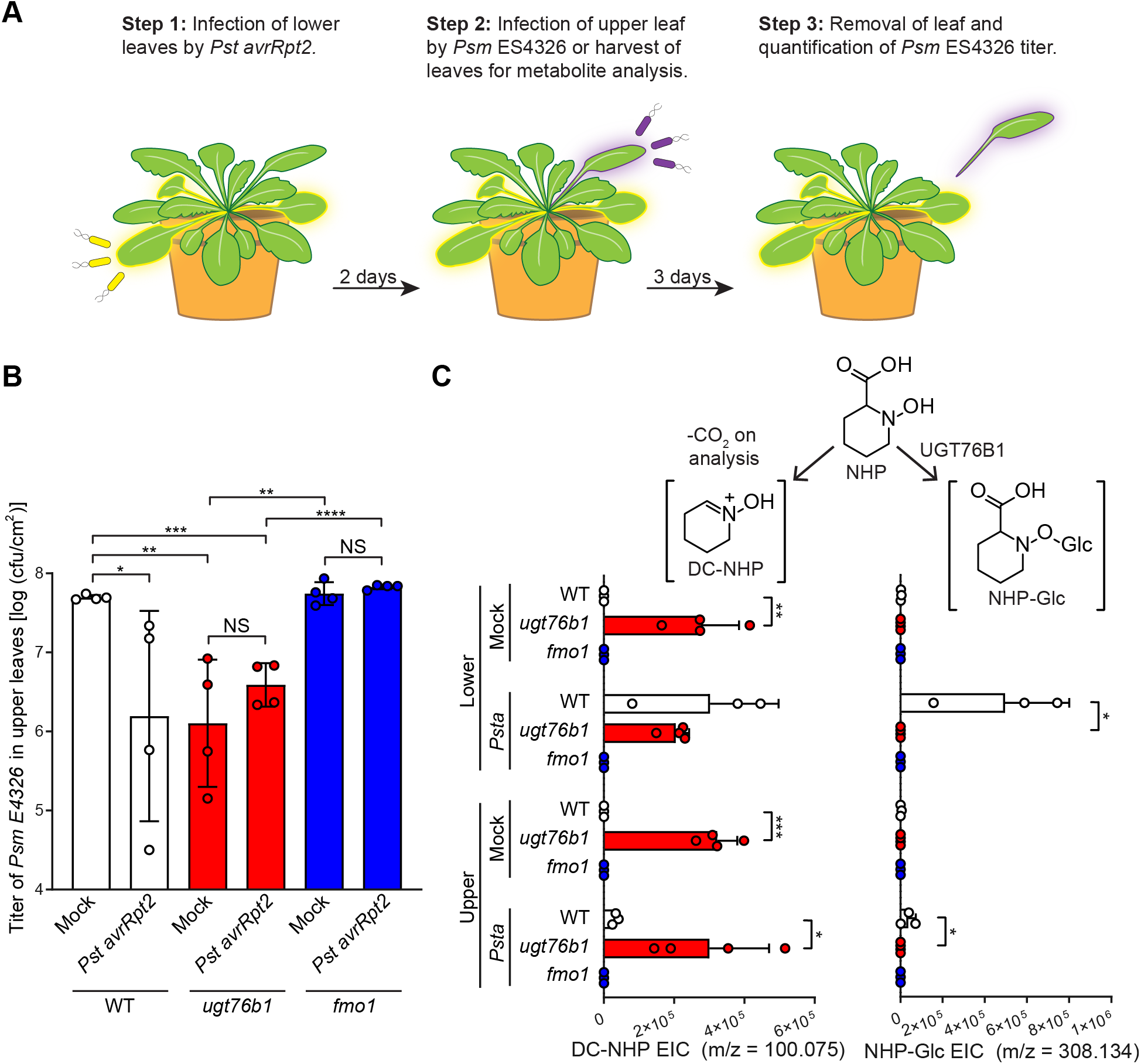
SAR assays in Arabidopsis WT, *ugt76b1* and *fmo1* plants. (A) Design of SAR assays in Arabidopsis. Three lower leaves (leaf number 5-7) of each plant were infiltrated with a 5 × 10^6^ cfu/ml suspension of *Pst avrRpt2* (*Psta*) (local infection) or 10 mM MgCl_2_ as a mock control. For bacterial growth assays in (B): two days after local infection, one upper leaf (leaf number 10) of each plant was challenged with 1 × 10^5^ cfu/ml suspension of *Psm* ES4326 (distal infection). Three days later, the disease symptoms of upper leaves were photographed and the titer of *Psm* ES4326 was determined. For metabolite analysis in (C): two days after local infection with *Pst avrRpt2*, the three lower infected leaves and three upper uninfected leaves (leaf numbers 8, 9, and 10) were harvested and separately pooled for metabolite analysis. (B) Titer of *Psm* ES4326 in upper, challenged leaves of WT (white bars), *ugt76b1* (red bars), and *fmo1* (blue bars) plants. Bars represent the mean ± SD (n = 4 independent biological replicates). Asterisks indicate a significant change in bacterial titer (one-tailed *t* test; \**P* < 0.05, \*\**P* < 0.01, \*\*\**P* < 0.001, \*\*\*\**P* < 0.0001, NS – not significant). The experiment was repeated three times with similar results. (C) Extracted ion abundances of DC-NHP (a degradation product of NHP) and NHP-Glc in methanolic tissue extracts from lower and upper leaves of WT (white bars), *ugt76b1* (red bars), and *fmo1* (blue bars) plants. Bars represent the means ± SD (*n* = 3 or 4 independent biological replicates). DC-NHP and NHP-Glc were measured using LC-MS. Values reported as zero indicate no detection of metabolites. Asterisks indicate a significant metabolite increase or decrease (one-tailed *t* test; \**P* < 0.05, \*\**P* < 0.01, \*\*\**P* < 0.001).

Upper leaves from WT plants initially treated with *Pst avrRpt2* harbored significantly less growth of *Psm* ES4326 than did WT plants treated with mock (Figure 4B), and developed fewer disease symptoms (e.g. bacterial speck and chlorosis; Supplemental Figure 5B), indicating the establishment of SAR. By contrast, SAR protection was abolished in *fmo1* plants (Figure 4B and Supplemental Figure 5B). Notably, the titers of *Psm* ES4326 in upper leaves of mock-treated *ugt76b1* plants were significantly lower than those of mock-treated WT plants. In addition, the titers of *Psm* ES4326 in the upper leaves of *ugt76b1* plants treated with mock and *Pst avrRpt2* were similar (Figure 4B), indicating that an initial pathogen infection was not required for disease resistance in the *ugt76B1* leaves. We also observed that all lower leaves (mock or *Pst avrRpt2*) of *ugt76b1* plants showed early senescence on the leaf margin (Supplemental Figure 5B), consistent with a prior report (von Saint Paul et al., 2011). Taken together, these findings indicate that mutation of *UGT76B1* leads to enhanced resistance regardless of an initial pathogen infection.

Based on our observations that *ugt76b1* seedlings have altered abundances of NHP, SA, and their glycosylated forms (Figure 3 and Supplementary Figure 4), we next determined the abundance of these metabolites using a modification of the SAR assay. We used the same experimental setup; however, we did not challenge with *Psm* ES4326. Instead, we harvested lower and upper leaves 2 days after mock or *Pst avrRpt2* treatment for metabolite analysis. We detected high background levels of Pip, SA, and SA-Glc in mock-treated *ugt76b1* plants, indicating that these plants are already primed with both NHP- and SA-related metabolites (Supplementary Figure 5C and D). Neither *fmo1* nor *ugt76b1* plants accumulated any NHP-Glc, while WT plants showed significant increases in both lower and upper leaves (Figure 4C), confirming the requirement for these two enzymes in the NHP-Glc biosynthetic pathway. As previously reported (von Saint Paul et al., 2011), *ugt76b1* plants contained significantly more SA-Glc than WT plants in mock conditions, suggesting that other UGTs are still able to generate SA-Glc at appreciable levels in this context (Supplementary Figure 5D). We did not directly detect any free NHP in this experiment, and we hypothesize that this is due to the instability of the molecule (Chen et al., 2018). The abundance of decarboxylated NHP (DC-NHP; which has been reported as a degradation product of NHP (Chen et al., 2018)) was significantly elevated in mock- and *Pst avrRpt2*-treated *ugt76b1* plants (Figure 4C), suggesting the constitutive accumulation of NHP and its subsequent degradation (either *in planta* or during the metabolite extraction process). Taken together, these results indicate that enhanced resistance in *ugt76b1* is associated with elevated abundance of NHP- and SA-related metabolites in uninduced conditions.

### Expression of UGT76B1 abolishes NHP-induced protection in tomato

The enhanced resistance exhibited by Arabidopsis *ugt76b1* mutants with significantly reduced levels of NHP-Glc suggested that the glycosylation of NHP reduces its bioactivity as a SAR signaling molecule. To explore this idea, we employed a transient SAR assay in tomato to study the phenotypic effect of increasing the relative abundance of NHP-Glc relative to NHP. In previous work, we established that transient expression of Arabidopsis ALD1 and FMO1 in tomato leaflets proximal to the main stem is sufficient to induce the production of NHP and inhibit the growth of *Pst* in infected distal leaflets (Holmes et al., 2019). Notably, altering NHP levels in tomato did not lead to the production of NHP-Glc (Holmes et al., 2019). Therefore, we reasoned that we could use this heterologous system as an experimental platform to study the role Arabidopsis UGT76B1 and NHP-Glc in SAR without significant contribution from native tomato UGTs.

We hypothesized that overexpression of Arabidopsis UGT76B1 with ALD1 and FMO1 would increase the ratio of NHP-Glc relative to NHP in proximal tomato leaflets and decrease the SAR response in distal leaflets infected with *Pst*. To test this, we infiltrated the two proximal leaflets of a fully expanded tomato leaf with *Agrobacteria* strains harboring *GFP* or *GFP* + *ALD1* + *FMO1* (Pathway), or Pathway + *UGT76B1* (Figure 5A). Two days post-infiltration, we harvested both proximal and distal leaflets for metabolite analysis (Figure 5A, C, and D). Proximal leaflets accumulated significantly less NHP and SA when expressing UGT76B1 alongside the NHP biosynthetic genes than did leaflets expressing NHP biosynthetic genes alone (Figure 5C and D). Conversely, these leaflets accumulated significantly more NHP-Glc and SA-Glc, suggesting direct conversion of the aglycones (Figure 5C and D). The only significant metabolic change that occurred in distal leaflets was an accumulation of free SA in leaves expressing only the NHP metabolic pathway enzymes (Supplemental Figure 6).

**Figure 5.**
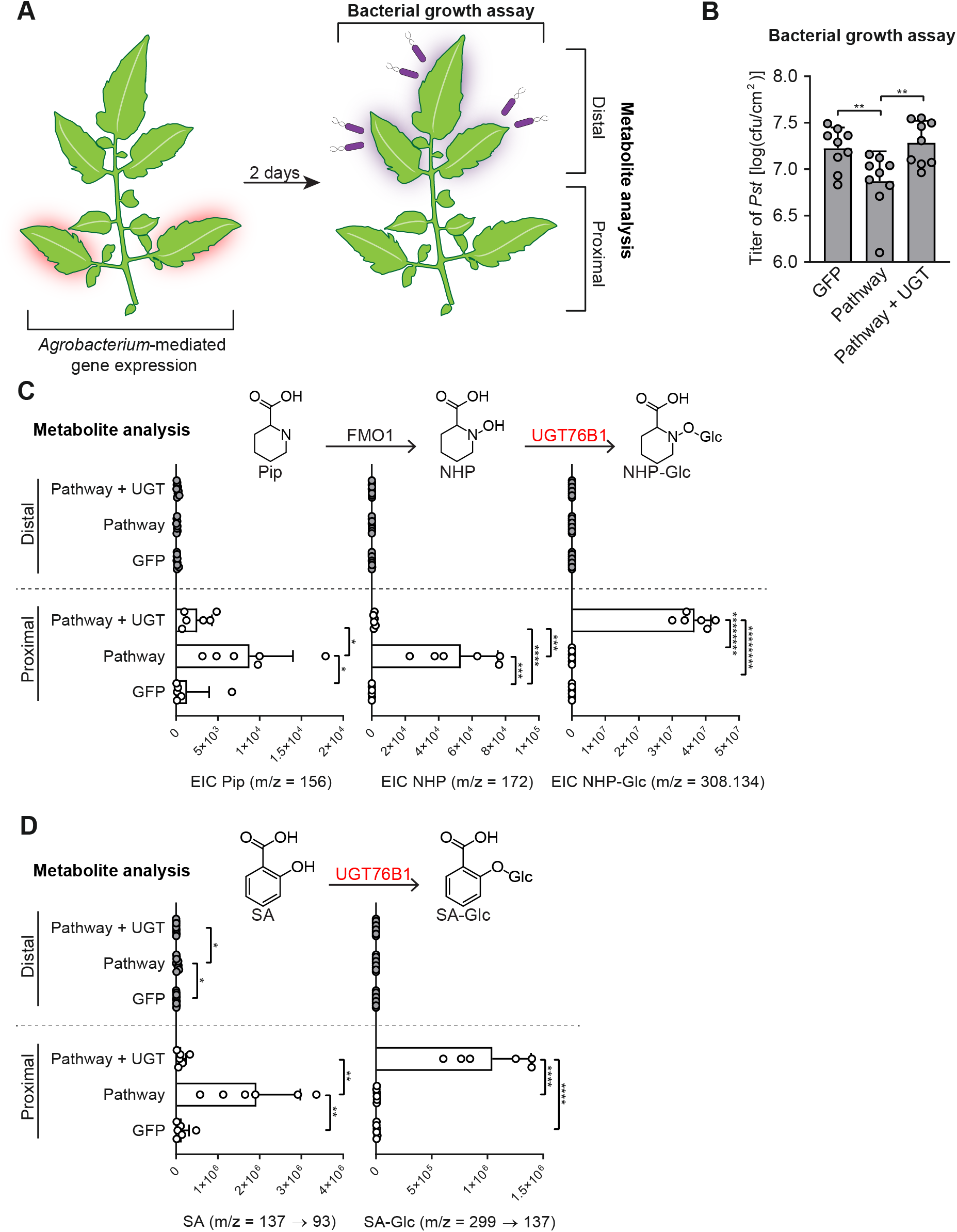
Transient expression of Arabidopsis *UGT76B1* with *ALD1* and *FMO1* in tomato leaves. (A) Design of transient SAR assays in tomato. Two leaflets of a tomato leaf proximal to the main stem (highlighted in red) were inoculated with *Agrobacteria* harboring *GFP* (GFP) or a combination of strains harboring *GFP* + Arabidopsis *ALD1* + Arabidopsis *FMO1* (Pathway) with out and with Arabidopsis *UGT76B1* (Pathway + UGT). For bacterial growth assay in (B): two days post infiltration with *Agrobacteria*, distal leaflets (highlighted in purple) were inoculated with a 1×10^5^ CFU/ml suspension of *Pst*. Four days post infiltration (dpi), distal leaves were harvested for quantification of *Pst* titers. For metabolite analysis in (C) and (D): two dpi with *Agrobacteria*, both proximal leaflets infiltrated with *Agrobacteria* and distal, untreated leaflets were harvested independently for analysis. (B) Titer of *Pst* in distal leaflets four dpi. Bars represent mean log cfu/cm^2^ ± SD (three leaflets each from n = 3 independent plants). Asterisks indicate a significant difference (one-tailed *t* test; \*\**P* < 0.01). (C) Abundances of Pip, NHP, and NHP-Glc in tomato leaflets expressing GFP, Pathway, and Pathway + UGT (white bars) and leaflets distal to those infiltrated with *Agrobacteria* (grey bars). Bars for proximal leaflets represent means ± SD (two leaflets each from n = 3 independent plants). Bars for distal leaflets represent means ± SD (three leaflets each from n = 3 independent plants). Pip and NHP were measured as TMS and 2-TMS derivatives respectively using GC-MS. NHP-Glc was measured using LC-MS. Values reported as zero indicate no detection of metabolites. Asterisks indicate a significant metabolite difference (one-tailed *t* test; \**P* < 0.05, \*\*\**P* < 0.001, \*\*\*\**P* < 0.0001, \*\*\*\*\*\*\**P* < 1×10^−6^, \*\*\*\*\*\*\*\**P* < 1×10^−7^). (D) Abundances of SA and SA-Glc in tomato leaflets expressing GFP, Pathway, and Pathway + UGT (white bars) and leaflets distal to those infiltrated with *Agrobacteria* (grey bars). Bars for proximal leaflets represent means ± SD (two leaflets each from n = 3 independent plants). Bars for distal leaflets represent means ± SD (three leaflets each from n = 3 independent plants). SA and SA-Glc were measured using LC-MS. Values reported as zero indicate no detection of metabolites. Asterisks indicate a significant metabolite difference (one-tailed *t* test; \*\**P* < 0.01, \*\*\*\**P* < 0.0001).

Using the same experimental design as for metabolite profiling, we inoculated two proximal tomato leaflets with *Agrobacteria* strains harboring *GFP* or *GFP* + *ALD1* + *FMO1* (Pathway), or Pathway + *UGT76B1* (Figure 5A). At 48 h post inoculation, we challenged three distal leaflets with a 1×10^5^ cfu/ml suspension of *Pst*. Consistent with a previous report (Holmes et al., 2019), transient expression of the NHP pathway in proximal leaflets resulted in significant protection against *Pst* in challenged distal leaflets when compared to that of transient expression of GFP alone (Figure 5B). Notably, this systemic resistance was compromised when UGT76B1 was overexpressed alongside the NHP pathway in proximal leaflets. Expressing UGT76B1 alone did not alter protection when compared to expressing GFP alone (Supplemental Figure 7). Together, these results demonstrate that overexpression of UGT76B1 is sufficient to convert NHP to NHP-Glc. Moreover, these data indicate that increasing the abundance of NHP-Glc is not sufficient to induce defense priming.

## Discussion

UGTs are a highly expanded class of biosynthetic enzymes in plants with 120 members in the Arabidopsis genome (Paquette et al., 2003). Characterized UGTs from Arabidopsis have diverse roles, including detoxification of xenobiotic substrates and regulation of active hormone levels. Substrate promiscuity is a feature of some plant UGTs, allowing them to conjugate diverse xenobiotic substrates (Osmani et al., 2009) while others have evolved to be far more specific, including the glycosyltransferases UGT74F1 and UGT74F2 which glycosylate SA in a regiospecific manner (George Thompson et al., 2017). The role of hormone-specific UGTs is often to generate inactive, yet stable storage forms that, in some cases, may be hydrolyzed back into active molecule (Westfall et al., 2013). Our results indicate that NHP-Glc is an inactive or less active derivative of NHP in Arabidopsis and tomato. It is unknown whether NHP-Glc can be enzymatically hydrolyzed back into NHP to reactivate immune signaling; however, little to no NHP-Glc accumulates in Arabidopsis plants in the absence of infection (Chen et al., 2018), which suggests that NHP biosynthesis is the primary mechanism to initiate NHP-dependent SAR signaling.

By mining publicly available Arabidopsis mRNA expression data, we found that the core NHP biosynthetic genes *ALD1* and *FMO1* appear to be tightly co-regulated. Even though *UGT76B1* is induced under many pathogen stress conditions, its expression is not as highly correlated with the core pathway genes across these same conditions (Supplemental Figure 8). We observed a similar phenomenon with known SA *UGTs* (Supplementary Figure 8). This suggests that differential expression of hormone modifying *UGTs* with different pathogen stressors may help coordinate dynamic immune responses. This highlights the importance of further investigation into the how transcriptional and/or posttranscriptional regulation of *UGT76B1* expression affects the abundance of bioactive metabolites during SAR. It has also been reported that *UGT76B1* is constitutively expressed in roots (von Saint Paul et al., 2011), suggesting that it may also play role in tissue-specific regulation of metabolite levels.

While NHP-Glc is the primary form of NHP detected in Arabidopsis and in the closely-related plant *Brassica rapa* (Chen et al., 2018; Holmes et al., 2019), we did not observe accumulation of NHP-Glc in *N. benthamiana* or tomato without ectopic expression of Arabidopsis *UGT76B1*. These data differ from patterns of accumulation observed for SA-Glc, which has been reported in diverse plant families, including the Solanaceae (Lee et al., 1995). We have shown that expression of UGT76B1 can inactivate NHP-related pathogen defense in tomato (Figure 5), which raises the question of how, or if, tomato can natively modulate abundance of NHP, the active SAR signal. Other compounds downstream of NHP have been reported during transient expression of NHP biosynthetic enzymes in *N. benthamiana* (Chen et al., 2018; Holmes et al., 2019), and these may represent distinct mechanisms that have evolved to modulate the abundance of active hormone during defense in other plant species.

Previous work has shown that *ugt76b1* plants have increased resistance to *Pst* and altered SA-dependent gene expression in local tissues (von Saint Paul et al., 2011). Our results reveal that *ugt76b1* plants have a basal level of resistance to *Psm* ES4326 infection equivalent to an SAR response induced in WT plants (Figure 4). It is possible that the increased availability of free NHP may be a driver of this phenotype in *ugt76b1*, as these plants have little to no ability to glycosylate NHP (Figure 3 and Supplemental Figure 4) and that NHP is known to be a potent modulator of defense (Chen et al., 2018; Hartmann et al., 2018).

Many aspects of plant defense are intimately intertwined. Complex regulatory mechanisms underlie responses to different pathogens (Glazebrook, 2005) and coordination of SAR (Shah et al., 2014). Vital components of the Arabidopsis signaling network include NHP and SA, which are both required to establish functional SAR (Klessig et al., 2018; Hartmann and Zeier, 2019). RNA sequencing has uncovered a large overlap between NHP- and SA-dependent gene regulation, but also the presence of SA-independent regulation in SAR (Bernsdorff et al., 2016; Hartmann et al., 2018; Hartmann and Zeier, 2019). Our biochemical studies now show that the enzyme previously known to glycosylate SA and ILA (von Saint Paul et al., 2011; Maksym et al., 2018) can also metabolize NHP (Supplemental Figure 4 and Supplemental Figure 5), further connecting these signaling molecules. It is possible that the true biological function of UGT76B1 is to glycosylate a set of small molecules and that all of them play distinct roles in defense.

Notably, our experiments in tomato provide additional evidence that NHP is a bioactive signaling molecule in SAR and reveal that glycosylation can be used to modulate this systemic response. Simply by expressing UGT76B1 alongside the NHP biosynthetic enzymes in tomato, the beneficial effect of producing NHP was abolished (Figure 5B). This finding provides insight into how plants regulate potent immune signals and may be critical for engineering approaches that seek to tune enhanced resistance in tomato. Efforts to improve resistance using synthetic chemicals has been challenging due to an inherent imbalance of plant defense and growth in the presence of inducers (Heil et al., 2000; Huot et al., 2014). Engineering immunity using synthetic approaches will need to address defense-yield tradeoffs plants naturally make to balance limited resources (Mauch et al., 2001; Ning et al., 2017). While NHP may be protective in the context of infection, constitutive expression would likely cause unintended growth defects, and any stable system would require inducible control of pathway enzymes and a mechanism to attenuate the signal in the absence of infection. We have shown that UGT76B1 can eliminate the NHP-dependent SAR signal in tomato (Figure 5), and this activity could be leveraged to engineer dynamic control over crop defense.

In closing, our results reveal that metabolism by the UDP-glycosyltransferase UGT76B1 plays critical role in modulating Arabidopsis immunity by glycosylating NHP, the key chemical initiator of SAR. We anticipate that the association of UGT76B1 with NHP signaling will more broadly contribute to our understanding of how plants use metabolic transformations of small plant signals to tune the dynamics, tissue specificity, and spatial regulation of defense responses.

## Materials and methods

### Gene expression and correlation analysis

Arabidopsis microarray datasets were obtained from the NASCArrays database (Craigon et al., 2004) (indexed experiments can be found at http://arabidopsis.info/affy/link_to_iplant.html). Log-scaled gene expression ratios were calculated from experiments 120, 122, 123, 167, 169, 330, 415, and 447 as previously (Rajniak et al., 2015). Pearson’s *r* correlation coefficient between genes was calculated from log_2_ normalized expression data from these microarray datasets.

### Plant materials and growth conditions

For seedling hydroponics experiments, *A. thaliana* ecotype Col-0 (WT), homozygous Syngenta Arabidopsis Insertion Library (SAIL) (Sessions et al., 2002) or T-DNA insertional line (SAIL_1171_A11; *ugt76b1-1*; Col-0 background) seeds were surface sterilized with 50% ethanol for 1 minute, 50% bleach for 10 min, washed 3 times in sterile water, and resuspended in 1x Murashige-Skoog (MS) medium with vitamins (PhytoTechnology Laboratories) (pH 5.7). Seeds were placed into 3 ml of MS medium + 5 g/l sucrose in wells of 6-well culture plates (5 seeds/well). Plates were sealed with micropore tape (3M), vernalized at 4°C for 48 h, and transferred to a growth chamber at 50% humidity, 22°C, and 100 μmol/m^2^/s photon flux on a 16-h/8-h day/night cycle. After 1 week, spent medium was removed and replaced with 3 ml of fresh MS medium + 5 g/l sucrose. Plants were elicited after an additional week of growth. For adult Arabidopsis experiments, Col-0, *fmo1-1* (SALK_026163; Col-0 background), and *ugt76b1* plants were grown in a growth chamber at 80% humidity, 22°C, and 100 μmol/m^2^/s photon flux on a 16-h/8-h day/night cycle. For tomato (*S. lycopersicum* cultivar VF36) experiments, plants were grown in a greenhouse (16-h/8-h day/night cycle, 25°-28°C) for 4-5 weeks. *N. benthamiana* plants were grown in soil on a growth shelf with a 16-h light cycle for 4 weeks prior to transient expression.

### Cloning of Arabidopsis UGT candidate genes

*Agrobacterium tumefaciens* GV3101 and C58C1 pCH32 strains harboring Arabidopsis *ALD1* and *FMO1* genes in the pEAQ-HT vector (Peyret and Lomonossoff, 2013) were constructed previously (Holmes et al., 2019). Arabidopsis UGT candidates were polymerase chain reaction (PCR)-amplified from Arabidopsis WT complementary DNA (cDNA) using gene-specific primers (Supplemental Table 1), cloned into pEAQ-HT between *AgeI* and *SmaI* cut sites using Gibson assembly, and transformed into *E. coli* 10-β. Sequence-confirmed plasmids were then transformed into *A. tumefaciens* GV3101 using heat shock. For creation of a His-tagged construct, Arabidopsis *UGT76B1* was PCR-amplified from WT cDNA using gene-specific primers (Supplemental Table 1), cloned into the pET24b vector under control of a T7 promoter, and transformed into *E. coli* BL21.

### Transient expression in N. benthamiana

*Agrobacteria* strains were grown on LB agar plates with appropriate antibiotics for 24 h. Cells were scraped from plates with an inoculation loop, washed three times with *Agrobacterium* induction medium [10 mM MES buffer, 10 mM MgCl_2_, and 150 μM acetosyringone (pH 5.7)], resuspended in *Agrobacterium* induction medium, and incubated at room temperature for 2 h with agitation. For screening of candidate UGTs, *Agrobacteria* harboring *ALD1*, *FMO1*, and respective *UGT* genes were combined in equal proportions with each at an OD_600_ of 0.1. In all cases, *Agrobacteria* harboring *GFP* was added to ensure an equal final OD_600_ of 0.6. These solutions were infiltrated into leaves of 4-week-old *N. benthamiana* plants using needleless syringes. Plants were incubated for 72 h on growth shelves on a 16-h light/8-h dark cycle prior to sample harvest.

### Sample harvest and derivatizations

For all metabolomics experiments, plant tissue was harvested, lyophilized to dryness, and homogenized using a ball mill (Retsch MM 400) at 25 Hz for 2 min. Single-well Arabidopsis hydroponics samples were resuspended in 500 μl of 80:20 MeOH:H_2_O and incubated at 4°C for 10 min. *N. benthamiana* and tomato samples were resuspended in 20 μl of 80:20 MeOH:H_2_O per mg dry tissue and incubated at 4°C for 10 min. The liquid fraction of each sample was split for LC-MS and GC-MS analysis respectively. Samples for GC-MS analysis were further derivatized with *N*-methyl-*N*-(trimethylsilyl)trifluoroacetamide (MSTFA) (Holmes et al., 2019). Samples were derivatized with trimethylsilyldiazomethane (TMSD) using previously established methods (Topolewska et al., 2015). Briefly, 200 μl dried methanolic extracts were resuspended in 125 μl, methanol, 50 μl toluene, and 50 μl 2M TMSD in hexane, incubated for 1 h at room temperature, dried under N2, and resuspended in 200 μl AcN + 0.1% formic acid for LC-MS analysis.

### LC-MS analysis

NHP, NHP-Glc and TMSD-derivatized NHP and NHP-Glc were measured using previously published methods on an Agilent 1260 HPLC coupled to an Agilent 6520 quadrupole time-of-flight electrospray ionization (Q-TOF ESI) mass spectrometer (Chen et al., 2018). For *in vitro* metabolomics experiments, SA-Glc and ILA-Glc were measured using the same parameters except in negative ionization mode. NHP-Glc and TMSD-derivatized NHP compounds were fragmented using a collision-induced dissociation energy (CID) of 10 V. TMSD-derivatized NHP-Glc was fragmented using a CID of 40 V. Extracted ion chromatogram (EIC) values were determined by extracting chromatograms with a 20 ppm error and integrating peak areas using MassHunter software (Agilent).

SA and SA-Glc were measured using an Agilent 1290 Infinity II UHPLC coupled to an Agilent 6470 triple quadrupole (QQQ) mass spectrometer. A 1.8 μm, 2.1 × 50 mm Zorbax RRHD Eclipse Plus C18 column was used for reverse phase chromatography with mobile phases of A [water with 0.1% formic acid (FA)] and B [acetonitrile (AcN) with 0.1% FA]. The following gradient was used for separation with a flow rate of 0.6 ml/min (percentages indicate percent buffer B): 0-0.2 min (5%), 0.2-4.2 min (5-95%), 4.2-5.2 min (95-100%). The MS was run in negative mode with the following parameters: gas temperature, 250C; gas flow rate, 12 l/min; nebulizer, 25 psig. SA was measured using monitored transitions with the following parameters: Precursor ion, 137.0239; product ions, 93 and 65.1; dwell, 150 ms; fragmentor voltage, 158 V; collision energy, 20 V and 32 V respectively, cell accelerator voltage, 4 V. SA-Glc was measured using monitored transitions with the following parameters: Precursor ion, 299.0767; product ions, 137 and 93; dwell, 150 ms; fragmentor voltage, 158 V; collision energy, 5 V and 20 V respectively, cell accelerator voltage, 4 V.

### GC-MS analysis

TMS-derivatized samples were measured for Pip and NHP using published methods on an Agilent 7820A gas chromatograph coupled to an Agilent 5977B mass spectrometer (Holmes et al., 2019).

### Bacterial strains and growth conditions

*Escherichia coli* strain 10-β, *Pseudomonas syringae* strains *pv. tomato* DC3000 (*Pst*), *pv. maculicola* ES4326 (*Psm* ES4326), and *pv. tomato* harboring the avirulence gene *avrRpt2* (*Pst avrRpt2*), and *Agrobacterium tumefaciens* strains GV3101 and C58C1 pCH32 were used in this study. *E. coli* strains were grown in lysogeny broth (LB) agar containing appropriate antibiotics at 37°C. *Pseudomonas* strains were grown at 28°C on nutrient yeast glycerol agar (NYGA) medium containing rifampicin (100 μg/ml). *Agrobacteria* strains were grown at 28°C on LB agar containing rifampicin (100 μg/ml), tetracycline (5 μg/ml), and kanamycin (50 μg/ml) for C58C1 pCH32 and gentamycin (100 μg/ml) and kanamycin (50 μg/ml) for GV3101.

### Elicitation methods

For hydroponics experiments, *Pst* was grown on LB agar plates at 30°C. A single colony was grown in liquid LB media to an OD_600_ of ~0.5, washed three times, and resuspended to an OD_600_ of 0.1 in MS media + 5 g/l sucrose. 30 μl of 1 M MgCl_2_ (mock), 100 mM NHP, 10 mM SA, or the *Pst* solution were used for elicitations.

### SAR assays in Arabidopsis

SAR bacterial growth assays were performed as described (Chen et al., 2018). 30-32-day-old Col-0, *fmo1*, and *ugt76b1* plants were used in this assay. Briefly, three lower leaves (leaf number 5-7) of each plant were infiltrated with 10 mM MgCl_2_ or a 5×10^6^ cfu/ml suspension of *Pst avrRpt2* in 10 mM MgCl_2_. Two days later, one upper leaf (leaf number 10) of each plant was inoculated with a 1×10^5^ cfu/ml suspension of *Psm ES4326*, and then plants were kept with a dome to maintain humidity. The disease symptoms of *Psm ES4326* infected upper leaves were photographed at 3 dpi, and then titer of *Psm ES4326* in these leaves was quantified by homogenizing leaves discs in 1 ml of 10 mM MgCl_2_, plating appropriate dilutions on NYGA medium with rifampicin (100 μg/ml). Plates were incubated at 28 °C for 2 days prior to counting bacterial colonies.

### Metabolic profiling of defense priming in Arabidopsis

Three lower leaves (leaf number 5-7) of 30-32-day-old Col-0, *fmo1* and *ugt76b1* Arabidopsis plants were infiltrated with 10 mM MgCl_2_ and a 5×10^6^ cfu/ml suspension of *Pst avrRpt2* in 10 mM MgCl_2_. Forty-eight h later, the three treated lower leaves and three untreated upper leaves (leaf number 8-10) were harvested, pooled, respectively, then frozen in liquid nitrogen for metabolic profiling by GC-MS, LC-MS, and triple quadrupole (QQQ)-MS analysis.

### Transient expression and SAR assays in tomato

Transient expression and SAR assays were performed as previously (Holmes et al., 2019). Briefly, combinations of *Agrobacterium* C58C1 pCH32 strains harboring combinations of *GFP*, *FMO1*, *ALD1*, and *UGT76B1* were infiltrated into two proximal (bottom) leaflets of the third and fourth compound leaves of 4-5-week old tomato plants for 48 h. For metabolic profiling, two proximal and three distal leaflets of the third compound leaf were harvested. For tomato, the three distal leaflets of the fourth compound leaves were inoculated with a 1×10^5^ cfu/ml suspension of *Pst*. Plants were incubated for four additional days, and then the titer of *Pst* was determined by plating serial dilutions (Holmes et al., 2019).

### In vitro assays

Crude protein was extracted from 80 mg of fresh tissue from *N. benthamiana* leaves transiently expressing GFP, UGT76B1, or UGT76B1-6xHis using the P-PER plant protein extraction kit (Pierce). Crude extracts of GFP and UGT76B1 were used directly in *in vitro* metabolomics assays. Protein concentrations were determined using a bicinchoninic acid assay kit (Pierce). All *in vitro* metabolomics assays with protein from *N. benthamiana* were performed in 200 μl reaction volumes at room temperature with the following concentration of reagents: 0.1 M Tris-HCl pH 7.5, 5 mM UDP-Glucose, 1 μg total protein, and 0.5 mM aglycone (NHP, SA, or ILA). Reactions were quenched by adding 50 μl reaction to 150 μl AcN.

*E. coli* BL21 strains harboring His-tagged UGT76B1 were grown overnight at 37°C in LB. Two ml of overnight culture was inoculated into 25 ml LB and cultures were grown at 37°C to an OD_600_ of 0.6. Cultures were then induced with 0.5 mM IPTG and grown for an additional 5 hours at 28°C. Cells were harvested and disrupted using an emulsiflex B15 (Avestin). Soluble fractions were enriched using gravity flow through Ni-NTA agarose resin and eluted with increasing concentrations of imidazole. Proteins were concentrated using 30 kDa centrifugal filters and buffer-exchanged into 50 mM Tris-HCl, pH 8 with 10% glycerol and kept at −80°C for long-term storage.

*In vitro* time course experiments were performed with 1 μM enriched *E. coli* UGT76B1-6xHis protein fraction in 200 μl reactions containing 0.1 M Tris-HCl pH 7.5, 0.5 mM UDP-Glc, and aglycone substrates at a concentration of 0.5 mM. Reactions were monitored as a time course at 5, 10, 30, and 60 min. Free UDP was measured as a proxy for reaction progress using the UDP-Glo enzyme assay kit (Promega) (Zegzouti et al., 2013)

## Accession numbers

The sequence data for this article can be found in the Arabidopsis Genome Initiative under the following accession numbers: UGT71B4 (AT4G15260), UGT73B2 (AT4G34135), UGT73B3 (AT4G34131), UGT73C5 (AT2G36800), UGT73D1 (AT3G53150), UGT74F2 (AT2G43820), UGT76B1 (AT3G11340), UGT76F2 (AT3g55700), UGT85A1 (AT1G22400), UGT85A7 (AT1G22340), UGT86A2 (AT2G28080), UGT89A2 (AT5G03490), UGT73C3 (AT2G36780), UGT92A1 (AT5G12890), UGT73B4 (AT2G15490), UGT87A2 (AT2G30140), UGT76E12 (AT3G46660), ALD1 (AT2G13810), and FMO1 (AT1G19250). Germplasm used in this study includes *fmo1-1* (SALK_026163) and *ugt76b1-1* (SAIL_1171_A11).

## List of supplemental materials

**Supplemental Figure 1**. mRNA expression profiles of candidate Arabidopsis *UGT* genes (Supports Figure 1).

**Supplemental Figure 2**. Arabidopsis UGT76B1 glycosylates NHP in *N. benthamiana* (Supports Figure 1).

**Supplemental Figure 3**. Trimethylsilyldiazomethane (TMSD) derivatization of NHP and NHP-Glc (Supports Figure 1).

**Supplemental Figure 4**. Metabolic profiling of Arabidopsis WT and *ugt76b1* mutant seedlings (Supports Figure 3).

**Supplemental Figure 5**. Metabolic profiling of Arabidopsis WT, *fmo1* and *ugt76b1* plants in SAR experiment (Supports Figure 4).

**Supplemental Figure 6**. Abundance of SA in distal leaflets of tomato during transient expression of NHP-Glc pathway genes (Supports Figure 5).

**Supplemental Figure 7**. Effect of transient expression of Arabidopsis UGT76B1 in tomato leaves for transient SAR analysis (Supports Figure 5).

**Supplemental Figure 8**. mRNA expression and coexpression analysis of SA-Glc and NHP-Glc biosynthetic genes in Arabidopsis obtained from publicly available microarray data.

**Supplemental Table 1**. Primers used in this study.

## Acknowledgements

We thank George Lomonossoff (John Innes Centre) for providing the pEAQ plasmid. We thank K. Smith and J. Foret (Stanford) for helpful discussions. This work was supported by the Howard Hughes Medical Institute (E.S.S.), an National Science Foundation Graduate Research Fellowship DGE-1656518 (to E.C.H.), National Science Foundation IOS-1555957 and Binational Science Foundation Grant 2011069 (to M.B.M.) and Ministry of Science and Technology of Taiwan-105-2917-I-564-093 (to Y.C.C.).

## Author Contributions

E.C.H., Y.C.C., E.S.S. and M.B.M designed the research; E.C.H. and Y.C.C. performed research; E.C.H., Y.C.C., E.S.S. and M.B.M. analyzed data; and E.C.H., Y.C.C., E.S.S. and M.B.M. wrote the paper.

---

**Supplemental Figure 1.**
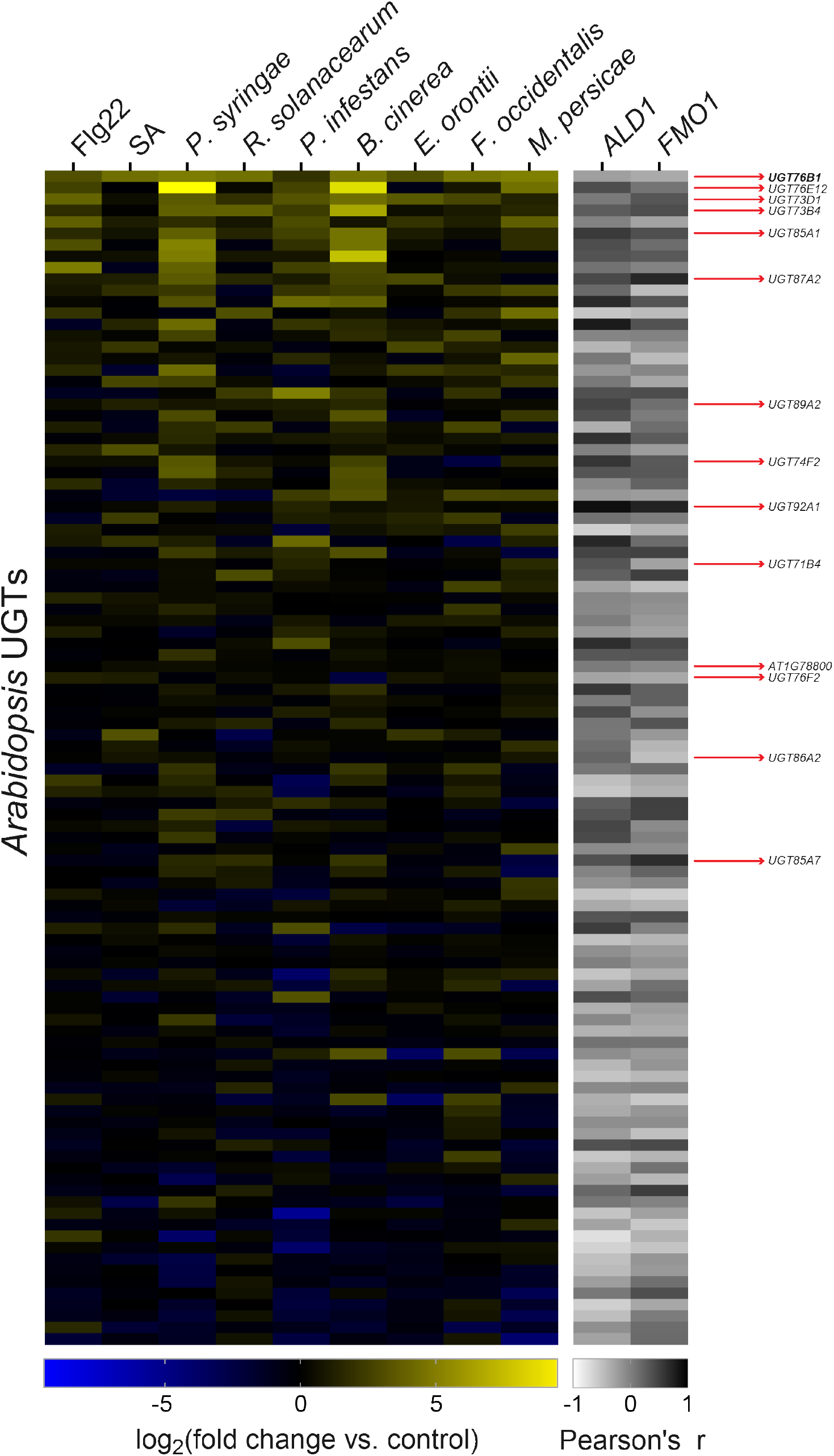
mRNA expression profiles of candidate Arabidopsis *UGT* genes (Supports Figure 1). Log transformed relative mRNA expression of 103 Arabidopsis UDP-dependent glycosyltransferases from publicly available microarray data. Log_2_(relative expression) is plotted on a linear gradient from −10 (blue) to 0 (black) to 10 (yellow). Pearson’s *r* correlation between the plotted expression patterns of the *UGT*s compared to respective NHP biosynthetic genes is plotted on a linear gradient from −1 (white) to 1 (black). *UGT*s are ordered by average relative expression across all biotic stress conditions: Flg22 (flagellin peptide), SA (salicylic acid hormone), bacterial pathogens (*Pseudomonas syringae* pv. *tomato* DC3000 and *Ralstonia solanacearum*), fungal/oomycete/ascomycete pathogens (*Botrytis cinerea*, *Phytophthora infestans*, and *Erysiphe orontii*), and insects/pests (*Frankliniella occidentalis* and *Myzus persicae*). *UGT*s tested in *N. benthamiana* during this study are indicated by red arrows. Four *UGT*s tested in *N. benthamiana* (*UGT73B2*, *UGT73B3*, *UGT73C3*, and *UGT73C5*) were not included in this expression analysis because they were not measured in the experiments analyzed.

**Supplemental Figure 2.**
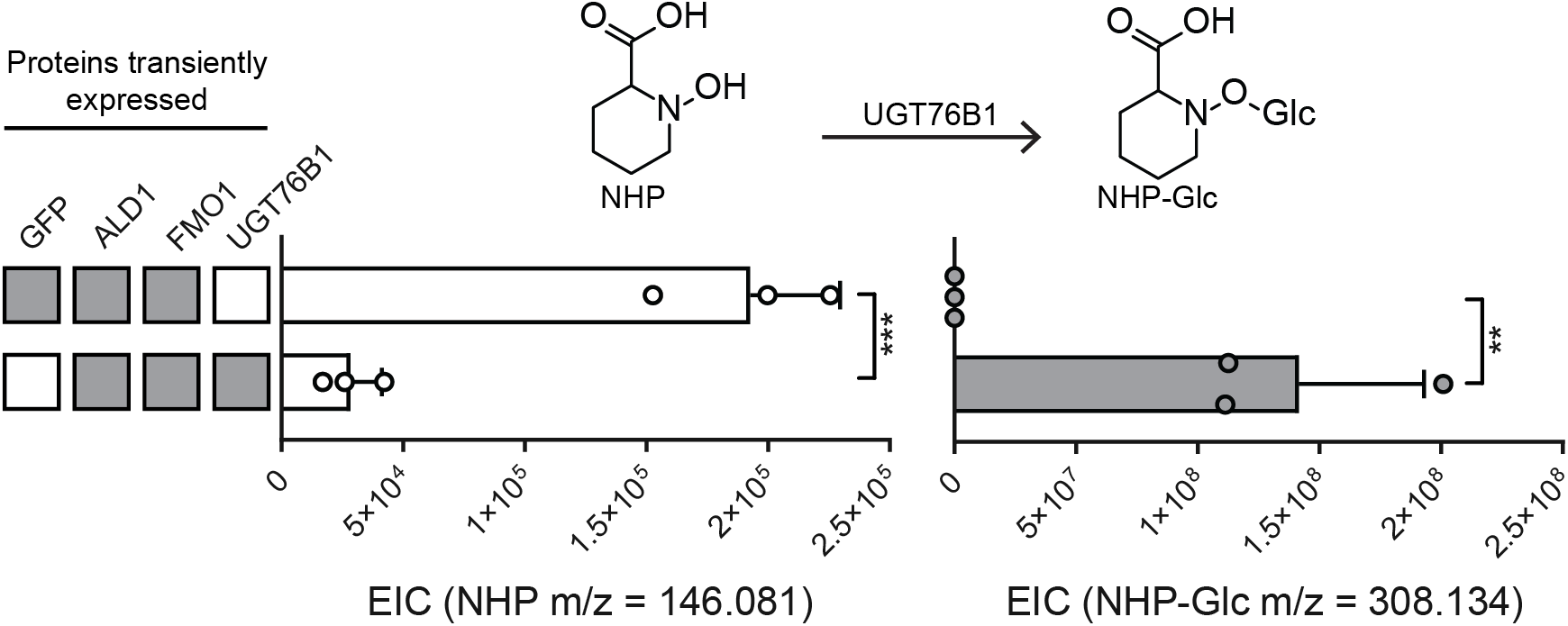
Arabidopsis UGT76B1 glycosylates NHP in *N. benthamiana* (Supports Figure 1). Abundances of NHP and NHP-Glc after transient expression of GFP + ALD1 + FMO1 and ALD1 + FMO1 + UGT76B1 in *N. benthamiana* leaves. Filled in grey boxes indicate an *Agrobacteria* strain including the respective gene was included in the experiment. Bars represent means ± SD (three independent biological replicates). NHP and NHP-Glc were measured using LC-MS. Values reported as zero indicate no detection of metabolites. Asterisks indicate a significant metabolite difference (one-tailed *t* test; \*\**P* < 0.01, \*\*\**P* < 0.001).

**Supplemental Figure 3.**
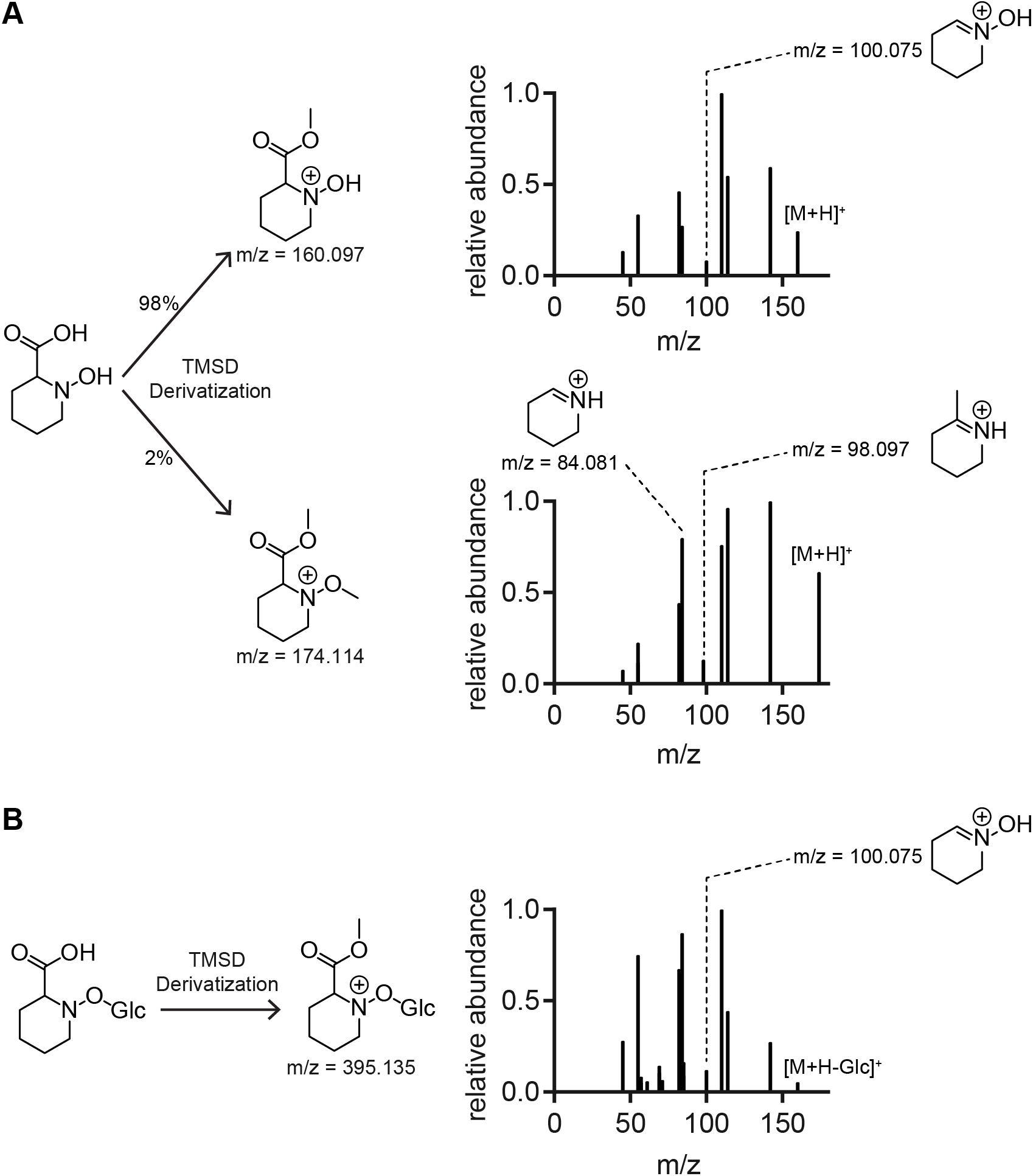
Trimethylsilyldiazomethane (TMSD) derivatization of NHP and NHP-Glc (Supports Figure 1). (A) TMSD derivatization of synthetic NHP yielded 98% singly methylated and 2% doubly methylated product by extracted ion chromatogram (EIC) quantification. The single methylation is hypothesized to occur on the acid based on MS/MS fragmentation and the presence of an m/z 100.075. MS/MS fragmentation for the doubly methylated product lacks m/z 100.075. (B) TMSD derivatization of extracts from *N. benthamiana* leaves expressing ALD1 + FMO1 + UGT76B1 led to a singly methylated NHP-Glc product. m/z 100.075 is present in the MS/MS fragmentation pattern, supporting methylation on the acid.

**Supplemental Figure 4.**
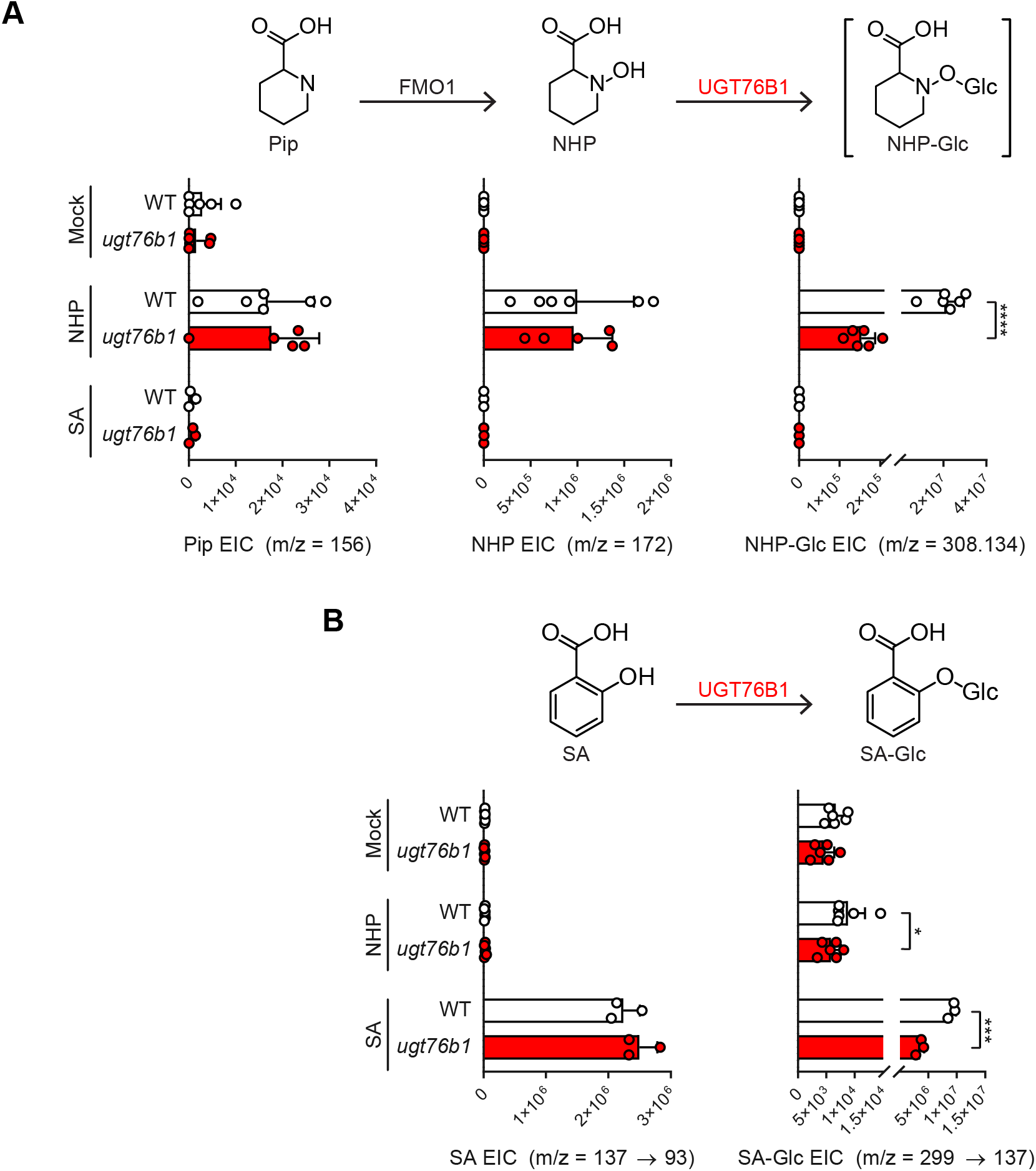
Metabolic profiling of Arabidopsis WT and *ugt76b1* mutant seedlings (Supports Figure 3). Arabidopsis WT (white bars) and *ugt76b1* (red bars) seedlings were grown hydroponically and treated with 1 mM MgCl_2_ (mock), 1 mM NHP, or 100 μM SA. Bars show abundances of NHP-related metabolites and SA-related metabolites (B) 24 h after treatment. Bars represent the means ± SD (*n* = 6 (mock and NHP treatments) or 3 (SA treatments) independent biological replicates). Pip and NHP were measured as TMS and 2-TMS derivatives respectively using GC-MS. NHP-Glc, SA, and SA-Glc were measured using LC-MS. Values reported as zero indicate no detection of metabolites. Asterisks indicate a significant metabolite decrease (one-tailed *t* test; \**P* < 0.05, \*\*\**P* < 0.001, \*\*\*\**P* < 0.0001).

**Supplemental Figure 5.**
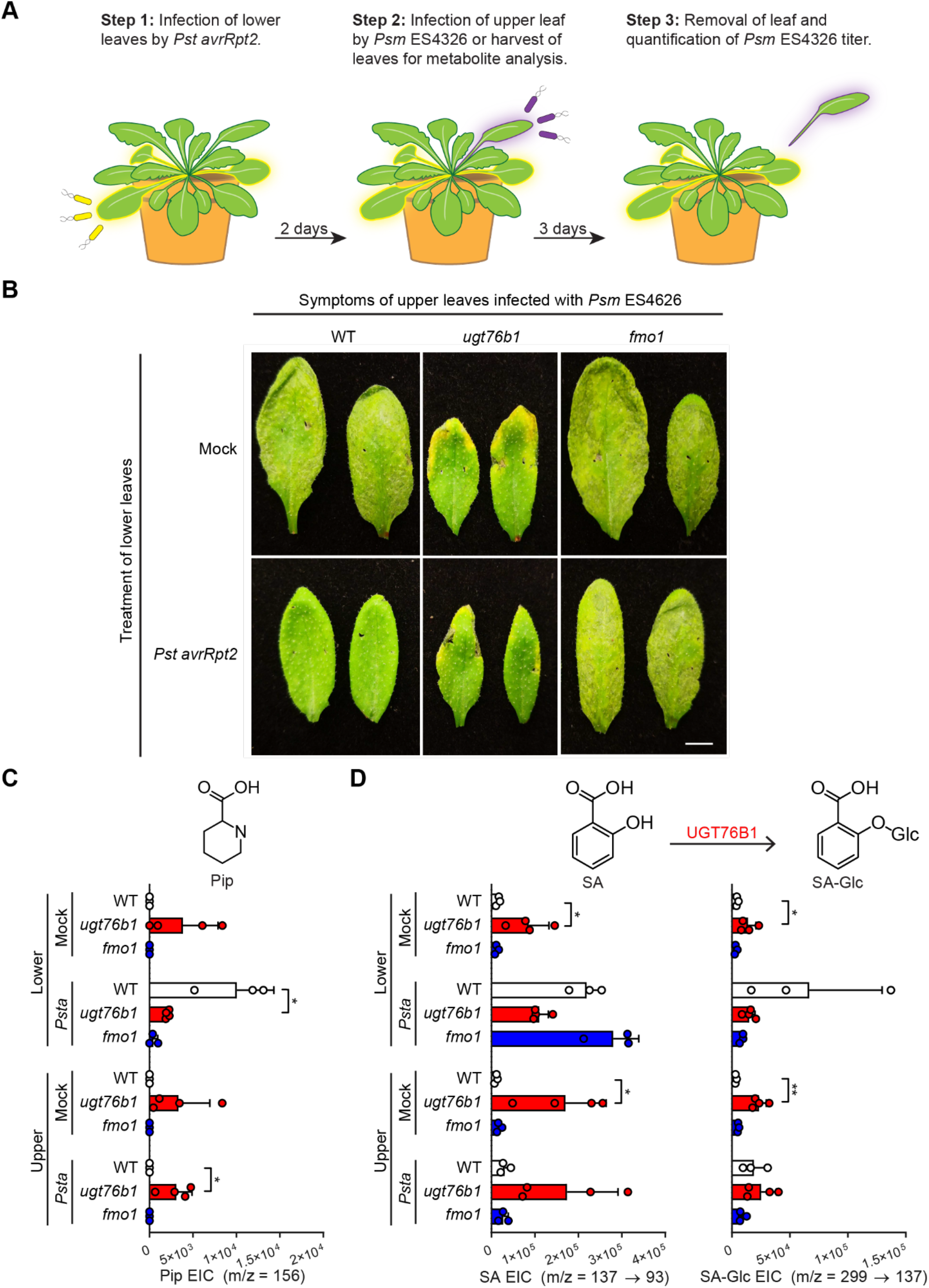
Metabolic profiling of Arabidopsis WT, *fmo1* and *ugt76b1* plants in SAR experiment (Supports Figure 4). (A) Design of SAR assays in Arabidopsis. Three lower leaves (leaf number 5-7) of each plant were infiltrated with a 5×10^6^ cfu/ml suspension of *Pst avrRpt2* (*Psta*) (local infection) or 10 mM MgCl_2_ as a mock control. For phenotype images in (B): two days after local infection, one upper leaf (leaf number 10) of each plant was challenged with 1× 0^5^ cfu/ml suspension of *Psm* ES4326 (distal infection). Three days later, the disease symptoms of upper leaves were photographed. For metabolite analysis in (C) and (D): two days after local infection with *Pst avrRpt2*, the three lower infected leaves and three upper uninfected leaves (leaf numbers 8, 9, and 10) were harvested and separately pooled for metabolite analysis. (B) Disease symptoms of two representative upper leaves inoculated with *Psm* ES4326 at 3 dpi. Scale bar = 0.5 cm. (C) Extracted ion abundances of Pip in methanolic tissue extracts from lower and upper leaves of WT (white bars), *ugt76b1* (red bars), and *fmo1* (blue bars) leaves. Bars represent the means ± SD (*n* = 3 or 4 independent biological replicates). Pip was measured as a TMS derivative using GC-MS. Values reported as zero indicate no detection of metabolites. Asterisks indicate a significant metabolite increase or decrease (one-tailed *t* test; \**P* < 0.05). (D) Extracted ion abundances of SA and SA-Glc in methanolic tissue extracts from lower and upper leaves of WT (white bars), *ugt76b1* (red bars), and *fmo1* (blue bars) plants. Bars represent the means ± SD (*n* = 3 or 4 independent biological replicates). SA and SA-Glc were measured using LC-MS. Values reported as zero indicate no detection of metabolites. Asterisks indicate a significant metabolite increase or decrease (one-tailed *t* test; \**P* < 0.05, \*\**P* < 0.01).

**Supplemental Figure 6.**
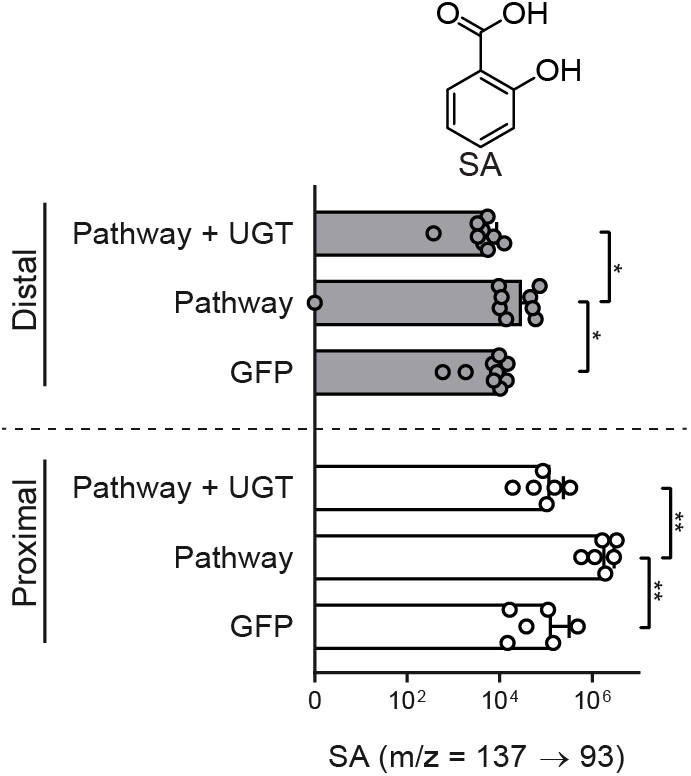
Abundance of SA in distal leaflets of tomato during transient expression of NHP-Glc pathway genes (Supports Figure 5). Abundance of SA in tomato proximal leaflets expressing GFP, Pathway (GFP + ALD1 + FMO1), and Pathway + UGT (white bars) and leaflets distal to those infiltrated with *Agrobacteria* (grey bars). Bars for proximal leaflets represent means ± SD (two leaflets each from n = 3 independent plants). Bars for distal leaflets represent means ± SD (three leaflets each from n = 3 independent plants). Values reported as zero indicate no detection of metabolites. SA was measured using LC-MS. Asterisks indicate a significant metabolite difference (one-tailed *t* test; \**P* < 0.05, \*\**P* < 0.01). Data are identical to that in Figure 5 with a different x-axis scale.

**Supplemental Figure 7.**
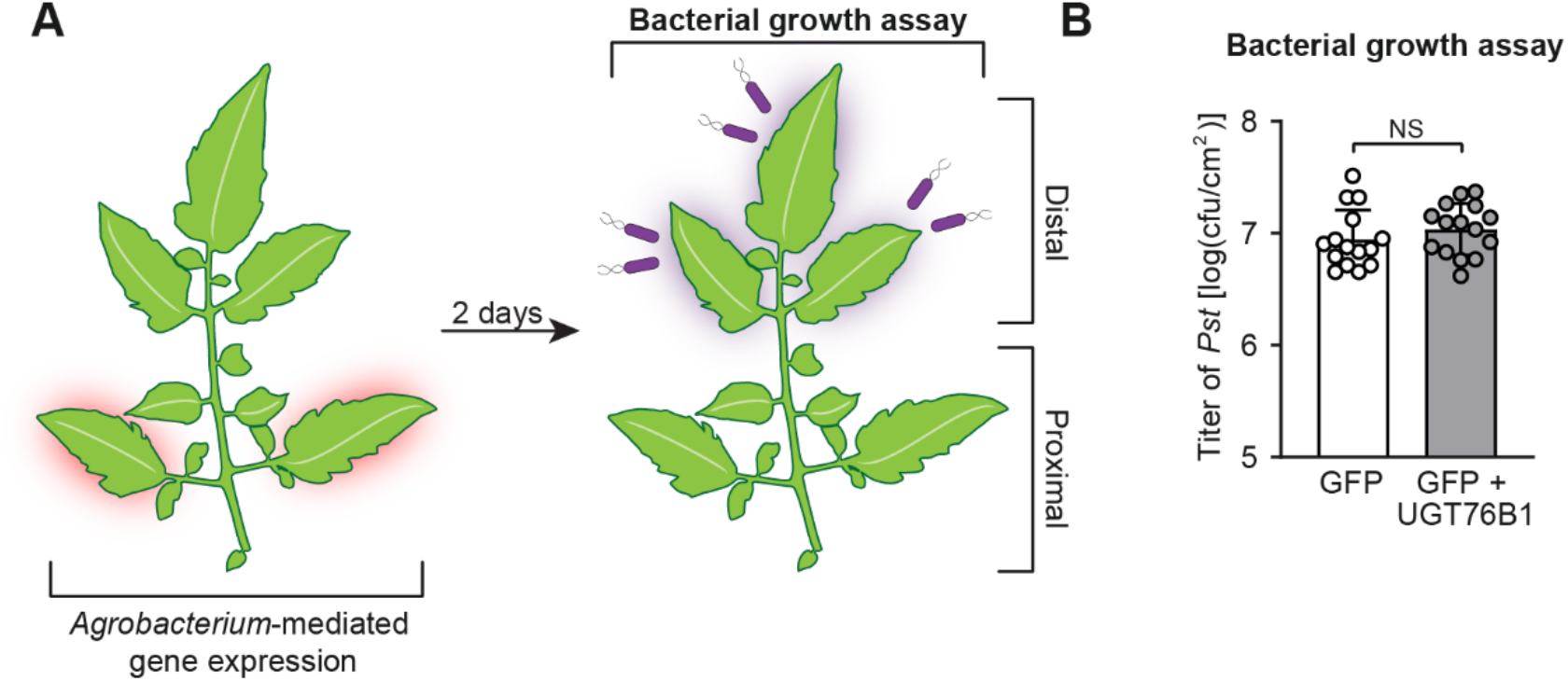
Effect of transient expression of Arabidopsis UGT76B1 in tomato leaves for transient SAR analysis (Supports Figure 5). (A) Design of transient SAR assays in tomato. Two leaflets of a tomato leaf proximal to the main stem (highlighted in red) were inoculated with *Agrobacteria* harboring *GFP* or *GFP* + Arabidopsis *UGT76B1*. For bacterial growth assay in (B): two days post infiltration with *Agrobacteria*, distal leaflets (highlighted in purple) were inoculated with a 1×10^5^ cfu/ml suspension of *Pst*. Four dpi, distal leaflets were harvested for quantification of *Pst* titers. (B) The titer of *Pst* in the distal leaflets was determined at 4 days post-inoculation. Bars represent means log(cfu/cm^2^) ± SD (three leaflets each from *n* = 5 independent plants). NS – not significant.

**Supplemental Figure 8.**
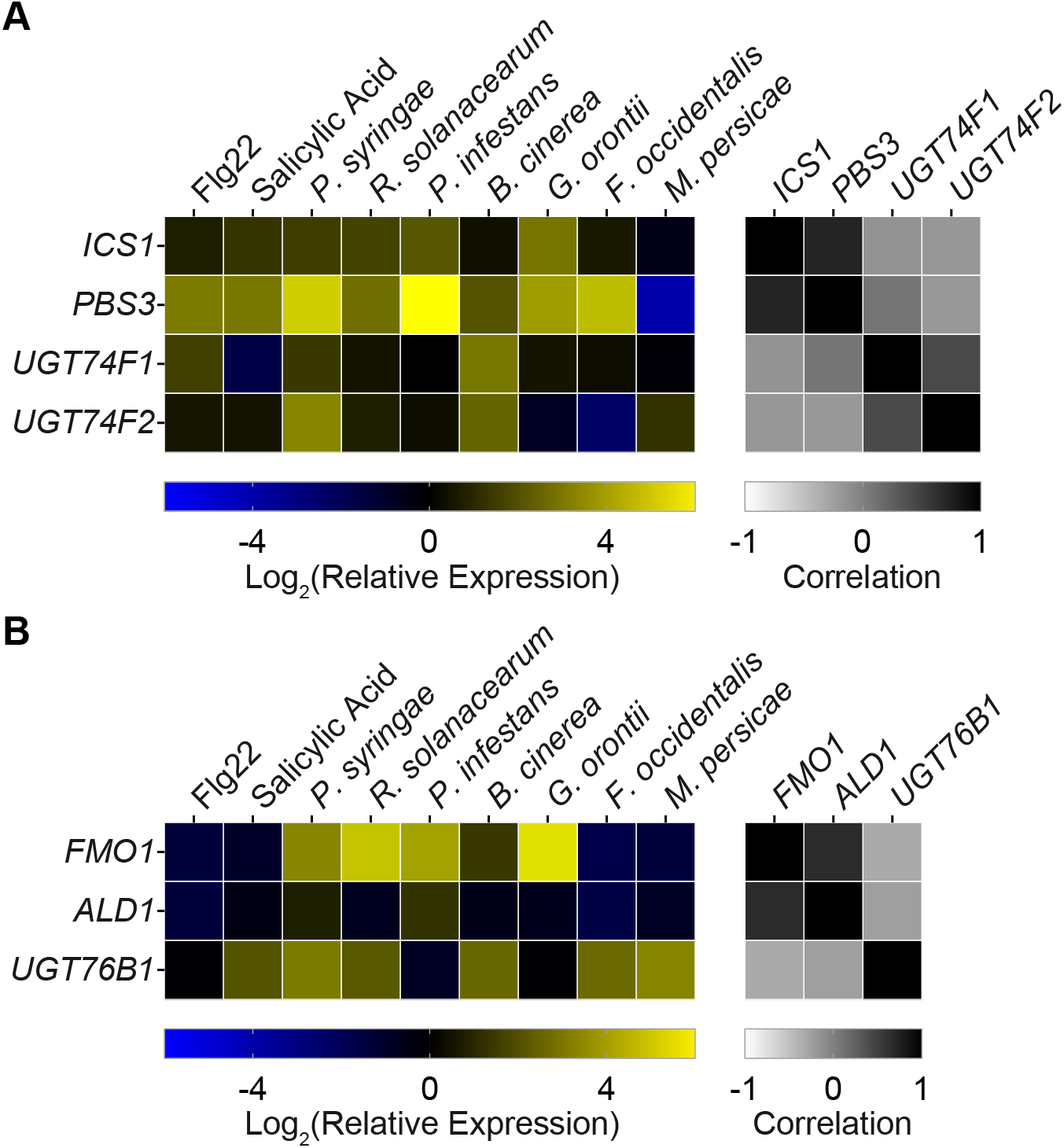
mRNA expression and coexpression analysis of SA-Glc and NHP-Glc biosynthetic genes in Arabidopsis obtained from publicly available microarray data. (A) Log transformed relative mRNA expression of SA biosynthetic genes (*ICS1; PBS3*) and glycosyltransferases (*UGT74F1; UGT74F2*) under biotic stress conditions. (B) Log transformed relative mRNA expression of NHP biosynthetic genes (*FMO1; ALD1*) and glycosyltransferase (*UGT76B1*) under biotic stress conditions. For both A and B, biotic treatments were: Flg22 (flagellin peptide), SA (salicylic acid hormone), bacterial pathogens (*Pseudomonas syringae* pv. *tomato* DC3000 and *Ralstonia solanacearum*), fungal/oomycete/ascomycetes pathogens (*Botrytis cinerea*, *Phytophthora infestans*, and *Erysiphe orontii*), and insects/pests (*Frankliniella occidentalis* and *Myzus persicae*). Log_2_(relative expression) is plotted on a linear gradient from −5 (blue) to 5 (yellow). Pearson’s *r* correlation between the plotted expression patterns of respective SA biosynthetic genes is plotted on a linear gradient from −1 (white) to 1 (black).

**Supplemental Table 1.**
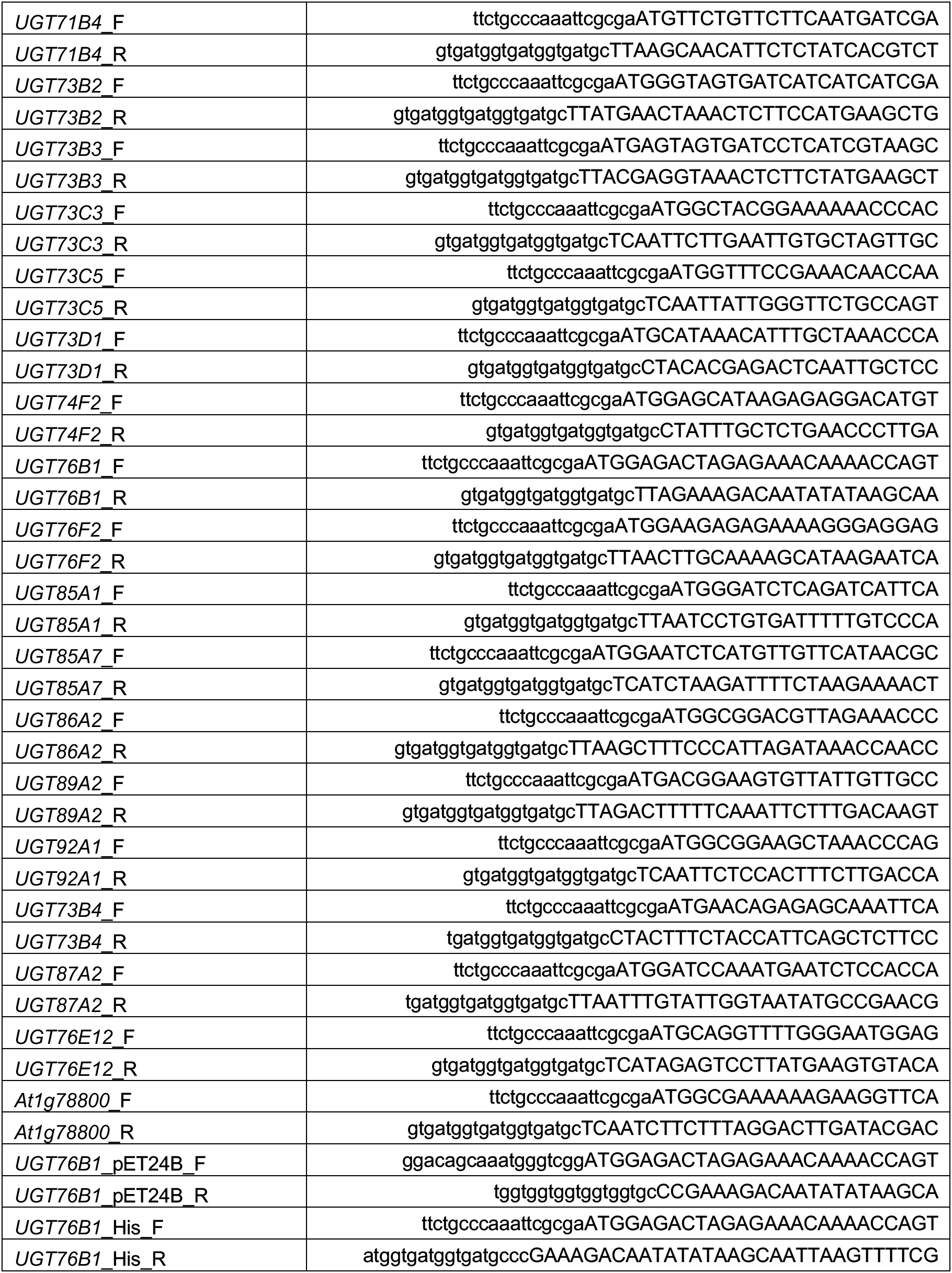
Primers used in this study. Lowercase letters indicate overlap with destination plasmid and uppercase letters indicate gene-specific sequence. F = Forward primer; R = Reverse primer. All sequences 5’-3’.

## Parsed Citations

Bauer, S., Mekonnen, D.W., Geist, B., Lange, B., Ghirardo, A., Zhang, W., and Schäffner, A.R. (2020). The isoleucic acid triad: distinct impacts on plant defense, root growth, and formation of reactive oxygen species. Journal of Experimental Botany, eraa160. Pubmed: Author and Title Google Scholar: Author Only Title Only Author and Title

Bernsdorff, F., Doring, A.C., Gruner, K., Schuck, S., Brautigam, A., and Zeier, J. (2016). Pipecolic acid orchestrates plant systemic acquired resistance and defense priming via salicylic acid-dependent and -independent pathways. Plant Cell 28, 102–129. Pubmed: Author and Title Google Scholar: Author Only Title Only Author and Title

Boachon, B., Gamir, J., Pastor, V., Erb, M., Dean, J.V., Flors, V., and Mauch-Mani, B. (2014). Role of two UDP-Glycosyltransferases from the L group of Arabidopsis in resistance against Pseudomonas syringae. European Journal of Plant Pathology 139, 707–720. Pubmed: Author and Title Google Scholar: Author Only Title Only Author and Title

Caarls, L., Elberse, J., Awwanah, M., Ludwig, N.R., de Vries, M., Zeilmaker, T., Van Wees, S.C.M., Schuurink, R.C., and Van den Ackerveken, G. (2017). Arabidopsis JASMONATE-INDUCED OXYGENASES down-regulate plant immunity byhydroxylation and inactivation of the hormone jasmonic acid. Proceedings of the National Academy of Sciences USA114, 6388. Pubmed: Author and Title Google Scholar: Author Only Title Only Author and Title

Chen, Y.-C., Holmes, E.C., Rajniak, J., Kim, J.-G., Tang, S., Fischer, C.R., Mudgett, M.B., and Sattely, E.S. (2018). N-hydroxy-pipecolic acid is a mobile metabolite that induces systemic disease resistance in Arabidopsis. Proceedings of the National Academy of Sciences USA, E4920–E4929. Pubmed: Author and Title Google Scholar: Author Only Title Only Author and Title

Choi, S., Cho, Y.-h., Kim, K., Matsui, M., Son, S.-H., Kim, S.-K., Fujioka, S., and Hwang, I. (2013). BAT1, a putative acyltransferase, modulates brassinosteroid levels in Arabidopsis. The Plant Journal 73, 380–391. Pubmed: Author and Title Google Scholar: Author Only Title Only Author and Title

Craigon, D.J., James, N., Okyere, J., Higgins, J., Jotham, J., and May, S. (2004). NASCArrays: a repositoryfor microarray data generated by NASC's transcriptomics service. Nucleic Acids Research 32, D575–D577. Pubmed: Author and Title Google Scholar: Author Only Title Only Author and Title

George Thompson, A.M., Iancu, C.V., Neet, K.E., Dean, J.V., and Choe, J.-y. (2017). Differences in salicylic acid glucose conjugations by UGT74F1 and UGT74F2 from Arabidopsis thaliana. Scientific Reports 7, 46629. Pubmed: Author and Title Google Scholar: Author Only Title Only Author and Title

Glazebrook, J. (2005). Contrasting mechanisms of defense against biotrophic and necrotrophic pathogens. Annual Review of Phytopathology 43, 205–227. Pubmed: Author and Title Google Scholar: Author Only Title Only Author and Title

Hartmann, M., and Zeier, J. (2018). l-lysine metabolism to N-hydroxypipecolic acid: an integral immune-activating pathwayin plants. The Plant Journal 96, 5–21. Pubmed: Author and Title Google Scholar: Author Only Title Only Author and Title

Hartmann, M., and Zeier, J. (2019). N-hydroxypipecolic acid and salicylic acid: a metabolic duo for systemic acquired resistance. Current Opinion in Plant Biology 50, 44–57. Pubmed: Author and Title Google Scholar: Author Only Title Only Author and Title

Hartmann, M., Zeier, T., Berndorff, F., Reichel-Deland, V., Kim, D., Hohmann, M., Scholten, N., Schuck, S., Brautigam, A., Holzel, T., Ganter, C., and Zeier, J. (2018). Flavin monooxygenase-generated N-hydroxypipecolic acid is a critical element of plant systemic immunity. Cell 173, 1–14. Pubmed: Author and Title Google Scholar: Author Only Title Only Author and Title

Heil, M., Hilpert, A., Kaiser, W., and Linsenmair, K.E. (2000). Reduced growth and seed set following chemical induction of pathogen defence: does systemic acquired resistance (SAR) incur allocation costs? Journal of Ecology 88, 645–654. Pubmed: Author and Title Google Scholar: Author Only Title Only Author and Title

Holmes, E.C., Chen, Y.-C., Sattely, E.S., and Mudgett, M.B. (2019). An engineered pathwayfor N-hydroxy-pipecolic acid synthesis enhances systemic acquired resistance in tomato. Science Signaling 12, eaay3066. Pubmed: Author and Title Google Scholar: Author Only Title Only Author and Title

Huot, B., Yao, J., Montgomery, B.L., and He, S.Y. (2014). Growth-defense tradeoffs in plants: Abalancing act to optimize fitness. Molecular Plant 7, 1267–1287. Pubmed: Author and Title Google Scholar: Author Only Title Only Author and Title

Kapila, J., De Rycke, R., Van Montagu, M., and Angenon, G. (1997). An Agrobacterium-mediated transient gene expression system for intact leaves. Plant Science 122, 101–108. Pubmed: Author and Title Google Scholar: Author Only Title Only Author and Title

Klessig, D.F., Choi, H.W., and Dempsey, D.M.A. (2018). Systemic acquired resistance and salicylic acid: Past, present, and future. Molecular Plant-Microbe Interactions 31, 871–888. Pubmed: Author and Title Google Scholar: Author Only Title Only Author and Title

Kühnel, E., Laffan, D.D.P., Lloyd-Jones, G.C., Martínez del Campo, T., Shepperson, I.R., and Slaughter, J.L. (2007). Mechanismof methyl esterification of carboxylic acids bytrimethylsilyldiazomethane. Angewandte Chemie International Edition 46, 7075–7078. Pubmed: Author and Title Google Scholar: Author Only Title Only Author and Title

Lee, H.I., León, J., and Raskin, I. (1995). Biosynthesis and metabolism of salicylic acid. Proceedings of the National Academy of Sciences USA92, 4076–4079. Pubmed: Author and Title Google Scholar: Author Only Title Only Author and Title

Lim, E.-K., Doucet, C.J., Li, Y., Elias, L., Worrall, D., Spencer, S.P., Ross, J., and Bowles, D.J. (2002). The activityof arabidopsis glycosyltransferases toward salicylic acid, 4-hydroxybenzoic acid, and other benzoates. Journal of Biological Chemistry 277, 586–592. Pubmed: Author and Title Google Scholar: Author Only Title Only Author and Title

Liu, P.-P., Yang, Y., Pichersky, E., and Klessig, D.F. (2009). Altering expression of benzoic acid/salicylic acid carboxyl methyltransferase 1 compromises systemic acquired resistance and PAMP-triggered immunity in Arabidopsis. Molecular Plant-Microbe Interactions 23, 82–90. Pubmed: Author and Title Google Scholar: Author Only Title Only Author and Title

Liu, Z., Yan, J.-P., Li, D.-K., Luo, Q., Yan, Q., Liu, Z.-B., Ye, L.-M., Wang, J.-M., Li, X.-F., and Yang, Y. (2015). UDP-Glucosyltransferase71C5, a major glucosyltransferase, mediates abscisic acid homeostasis in Arabidopsis. Plant Physiology 167, 1659. Pubmed: Author and Title Google Scholar: Author Only Title Only Author and Title

Maksym, R.P., Ghirardo, A., Zhang, W., von Saint Paul, V., Lange, B., Geist, B., Hajirezaei, M.-R., Schnitzler, J.-P., and Schäffner, A.R. (2018). The defense-related isoleucic acid differentiallyaccumulates in Arabidopsis among branched-chain amino acid-related 2-hydroxy carboxylic acids. Frontiers in Plant Science 9, 766. Pubmed: Author and Title Google Scholar: Author Only Title Only Author and Title

Mauch, F., Mauch-Mani, B., Gaille, C., Kull, B., Haas, D., and Reimmann, C. (2001). Manipulation of salicylate content in Arabidopsis thaliana bythe expression of an engineered bacterial salicylate synthase. The Plant Journal 25, 67–77. Pubmed: Author and Title Google Scholar: Author Only Title Only Author and Title

Nakamura, Y., Mithöfer, A., Kombrink, E., Boland, W., Hamamoto, S., Uozumi, N., Tohma, K., and Ueda, M. (2011). 12-Hydroxyjasmonic acid glucoside is a COI1-JAZ-independent activator of leaf-closing movement in Samanea saman. Plant Physiology 155, 1226. Pubmed: Author and Title Google Scholar: Author Only Title Only Author and Title

Nakazawa, M., Yabe, N., Ichikawa, T., Yamamoto, Y.Y., Yoshizumi, T., Hasunuma, K., and Matsui, M. (2001). DFL1, an auxin-responsive GH3 gene homologue, negativelyregulates shoot cell elongation and lateral root formation, and positivelyregulates the light response of hypocotyl length. The Plant Journal 25, 213–221. Pubmed: Author and Title Google Scholar: Author Only Title Only Author and Title

Navarova, H., Bernsdorff, F., Doring, A.C., and Zeier, J. (2012). Pipecolic acid, an endogenous mediator of defense amplification and priming, is a critical regulator of inducible plant immunity. The Plant Cell 24, 5123–5141. Pubmed: Author and Title Google Scholar: Author Only Title Only Author and Title

Ning, Y., Liu, W., and Wang, G.-L. (2017). Balancing immunity and yield in crop plants. Trends in Plant Science 22, 1069–1079. Pubmed: Author and Title Google Scholar: Author Only Title Only Author and Title

Noutoshi, Y., Okazaki, M., Kida, T., Nishina, Y., Morishita, Y., Ogawa, T., Suzuki, H., Shibata, D., Jikumaru, Y., Hanada, A., Kamiya, Y., and Shirasu, K. (2012). Novel plant immune-priming compounds identified via high-throughput chemical screening target salicylic acid glucosyltransferases in Arabidopsis. The Plant Cell 24, 3795. Pubmed: Author and Title Google Scholar: Author Only Title Only Author and Title

Osmani, S.A., Bak, S., and Møller, B.L. (2009). Substrate specificityof plant UDP-dependent glycosyltransferases predicted fromcrystal structures and homology modeling. Phytochemistry 70, 325–347. Pubmed: Author and Title Google Scholar: Author Only Title Only Author and Title

Paquette, S., Møller, B.L., and Bak, S. (2003). On the origin of family 1 plant glycosyltransferases. Phytochemistry 62, 399–413. Pubmed: Author and Title Google Scholar: Author Only Title Only Author and Title

Peyret, H., and Lomonossoff, G.P. (2013). The pEAQ vector series: the easyand quick wayto produce recombinant proteins in plants. Plant Molecular Biology 83, 51–58. Pubmed: Author and Title Google Scholar: Author Only Title Only Author and Title

Piotrowska, A., and Bajguz, A. (2011). Conjugates of abscisic acid, brassinosteroids, ethylene, gibberellins, and jasmonates. Phytochemistry 72, 2097–2112. Pubmed: Author and Title Google Scholar: Author Only Title Only Author and Title

Rajniak, J., Barco, B., Clay, N.K., and Sattely, E.S. (2015). Anew cyanogenic metabolite in Arabidopsis required for inducible pathogen defence. Nature 525, 376–379. Pubmed: Author and Title Google Scholar: Author Only Title Only Author and Title

Sessions, A., Burke, E., Presting, G., Aux, G., McElver, J., Patton, D., Dietrich, B., Ho, P., Bacwaden, J., Ko, C., Clarke, J.D., Cotton, D., Bullis, D., Snell, J., Miguel, T., Hutchison, D., Kimmerly, B., Mitzel, T., Katagiri, F., Glazebrook, J., Law, M., and Goff, S.A. (2002). Ahigh-throughput Arabidopsis reverse genetics system. The Plant Cell 14, 2985. Pubmed: Author and Title Google Scholar: Author Only Title Only Author and Title

Shah, J., Chaturvedi, R., Chowdhury, Z., Venables, B., and Petros, R.A. (2014). Signaling bysmall metabolites in systemic acquired resistance. The Plant Journal 79, 645–658. Pubmed: Author and Title Google Scholar: Author Only Title Only Author and Title

Smirnova, E., Marquis, V., Poirier, L., Aubert, Y., Zumsteg, J., Ménard, R., Miesch, L., and Heitz, T. (2017). Jasmonic acid oxidase 2 hydroxylates jasmonic acid and represses basal defense and resistance responses against Botrytis cinerea infection. Molecular Plant 10, 1159–1173. Pubmed: Author and Title Google Scholar: Author Only Title Only Author and Title

Staswick, P.E., and Tiryaki, I. (2004). The oxylipin signal jasmonic acid is activated byan enzyme that conjugates it to isoleucine in Arabidopsis. The Plant Cell 16, 2117. Pubmed: Author and Title Google Scholar: Author Only Title Only Author and Title

Staswick, P.E., Serban, B., Rowe, M., Tiryaki, I., Maldonado, M.T., Maldonado, M.C., and Suza, W. (2005). Characterization of an Arabidopsis enzyme family that conjugates amino acids to indole-3-acetic acid. The Plant Cell 17, 616. Pubmed: Author and Title Google Scholar: Author Only Title Only Author and Title

Takase, T., Nakazawa, M., Ishikawa, A., Kawashima, M., Ichikawa, T., Takahashi, N., Shimada, H., Manabe, K., and Matsui, M. (2004). ydk1-D, an auxin-responsive GH3 mutant that is involved in hypocotyl and root elongation. The Plant Journal 37, 471–483. Pubmed: Author and Title Google Scholar: Author Only Title Only Author and Title

Topolewska, A., Czarnowska, K., Haliński, Ł.P., and Stepnowski, P. (2015). Evaluation of four derivatization methods for the analysis of fattyacids fromgreen leafyvegetables bygas chromatography. Journal of Chromatography B 990, 150–157. Pubmed: Author and Title Google Scholar: Author Only Title Only Author and Title

von Saint Paul, V., Zhang, W., Kanawati, B., Geist, B., Faus-Keßler, T., Schmitt-Kopplin, P., and Schäffner, A.R. (2011). The Arabidopsis glucosyltransferase UGT76B1 conjugates isoleucic acid and modulates plant defense and senescence. The Plant Cell 23, 4124. Pubmed: Author and Title Google Scholar: Author Only Title Only Author and Title

Wang, B., Jin, S.-H., Hu, H.-Q., Sun, Y.-G., Wang, Y.-W., Han, P., and Hou, B.-K. (2012). UGT87A2, an Arabidopsis glycosyltransferase, regulates flowering time via FLOWERING LOCUS C. New Phytologist 194, 666–675. Pubmed: Author and Title Google Scholar: Author Only Title Only Author and Title

Wasternack, C., and Hause, B. (2013). Jasmonates: biosynthesis, perception, signal transduction and action in plant stress response, growth and development. An update to the 2007 review in Annals of Botany. Annals of Botany 111, 1021–1058. Pubmed: Author and Title Google Scholar: Author Only Title Only Author and Title

Westfall, C.S., Muehler, A.M., and Jez, J.M. (2013). Enzyme action in the regulation of plant hormone responses. Journal of Biological Chemistry 288, 19304–19311. Pubmed: Author and Title Google Scholar: Author Only Title Only Author and Title

Zegzouti, H., Engel, L., Vidugiris, G., and Goueli, S. (2013). Detection of glycosyltransferase activities with homogenous bioluminescent UDP detection assay. Glycobiology 23, 1340–1341. Pubmed: Author and Title Google Scholar: Author Only Title Only Author and Title

